# Heterologous betacoronavirus spike immunization in nonhuman primates elicits cross-reactive antibodies that neutralize both sarbeco- and merbecoviruses

**DOI:** 10.64898/2026.01.22.700386

**Authors:** Katharina Dueker, Tazio Capozzola, Ziqi Feng, Ryan N. Lin, Jonathan Hurtado, Sandhya Bangaru, Meng Yuan, Nathan Beutler, Elijah Garcia, Wan-ting He, Sean Callaghan, Gabriel Avillion, Lina Vo, Xuduo Li, Jonathan L. Torres, Rami Musharrafieh, Ge Song, Nitesh Mishra, Pragati Sharma, Peter Yong, Fabio Anzanello, Kasia Kaczmarek-Michaels, Elana Ben-Akiva, Murillo Silva, Mariane Melo, Muzamil Makhdoomi, Ethan Westfall-Gomez, William Rinaldi, Melissa Ferguson, Yana Safonova, Shane Crotty, Darrell J. Irvine, Thomas Rogers, Andrew B. Ward, Bryan Briney, Ian A. Wilson, Dennis R. Burton, Raiees Andrabi

**Affiliations:** Department of Immunology and Microbiology, The Scripps Research Institute, La Jolla, CA 92037, USA; IAVI Neutralizing Antibody Center, The Scripps Research Institute, La Jolla, CA 92037, USA; Consortium for HIV/AIDS Vaccine Development (CHAVD), The Scripps Research Institute, La Jolla, CA 92037, USA; Department of Medicine, University of Pennsylvania, Philadelphia, PA 19104, USA; Department of Integrative Structural and Computational Biology, The Scripps Research Institute, La Jolla, CA 92037, USA; Koch Institute for Integrative Cancer Research, Massachusetts Institute of Technology, Cambridge, MA 02139, USA; Howard Hughes Medical Institute, Chevy Chase, MD 20815, USA; Alpha Genesis, Yemassee, SC 29945, USA; Computer Science and Engineering Department, Huck Institutes of the Life Sciences, Penn State University, State College, PA 16801, USA; Division of Infectious Diseases and Global Public Health, Department of Medicine, University of California, San Diego (UCSD), La Jolla, CA 92037, USA; Center for Infectious Disease and Vaccine Research, La Jolla Institute for Immunology, La Jolla, CA 92037, USA; Department of Biological Engineering, Massachusetts Institute of Technology, Cambridge, MA 02139, USA; Skaggs Institute for Chemical Biology, The Scripps Research Institute, La Jolla, CA 92037, USA; Ragon Institute of Massachusetts General Hospital, Massachusetts Institute of Technology, and Harvard University, Cambridge, MA 02139, USA

## Abstract

In anticipation of future coronavirus (CoV) pandemics, developing vaccines that elicit broadly neutralizing antibodies (bnAbs) against diverse CoVs is critical. Here, we vaccinated rhesus macaques with SARS-CoV-2 spike (S)-protein, then boosted with heterologous β-CoV S-proteins to focus responses to common conserved S2 bnAb epitopes. Initial SARS-CoV-2 priming elicited receptor-binding domain (RBD)-focused responses, while MERS-CoV boosting redirected responses toward the S2 region, including the stem-helix bnAb site. Although S2-directed serum cross-neutralization was undetectable and most isolated cross-reactive monoclonal antibodies (mAbs) targeted non-neutralizing epitopes, two S2 stem-helix mAbs were identified from memory B cells. These bnAbs neutralized diverse sarbeco- and merbecoviruses, including MERS-CoV, and conferred robust *in vivo* protection against SARS-CoV-2 challenge. Structural studies revealed that these macaque bnAbs closely mimic human S2-stem bnAbs induced by infection. These findings provide proof-of-principle for vaccination strategies that elicit broadly protective β-coronavirus responses and highlight non-human primates as a translational model for evaluating S2-targeted immunogens.

## Introduction

In late 2019, the emergence of severe acute respiratory syndrome coronavirus-2 (SARS-CoV-2) caused the coronavirus disease 2019 (COVID-19) pandemic, leading to ∼7.0 million deaths worldwide and severely impacting the global economy (Who coronavirus (covid-19) dashboard). Taking into account the SARS-CoV-1 epidemic in 2002-2003 (Cherry and Krogstad, 2004), and the MERS-CoV outbreak in 2012 (Ramadan and Shaib, 2019), the SARS-CoV-2 pandemic in 2019 underscored yet again the tremendous threat that human coronaviruses (hCoVs) pose to human health. While the development and distribution of mRNA-based vaccines against SARS-CoV-2; BT162b2 (Polack et al., 2020) and mRNA-1273 (Baden et al., 2020) were a great success in reducing infection risk, symptom severity, and mortality, the extensive immune evasion observed in emerging SARS-CoV-2 variants of concern (VOCs) has since notably reduced the efficacy of the original wild-type vaccines (Collier et al., 2021; Han and Ye, 2022; Harvey et al., 2021; Krause et al., 2021; Paul et al., 2023; Wang et al., 2021b) and has required updates to COVID vaccines to include Omicron variants. The continuous emergence of immune evasive SARS-CoV-2 VOCs and the threat of possible future spillover events of coronaviruses suggest the need for vaccines that are insensitive to new and future emerging SARS-CoV-2 VOCs and which can lead to broad protection against β-coronaviruses.

Several pan-coronavirus immunization strategies have shown promise in preclinical animal models. Nanoparticle formulations displaying full-length spike protein (Hutchinson et al., 2023; Joyce et al., 2021; Joyce et al., 2022; Martinez et al., 2021), the receptor-binding domain (RBD) (Joyce *et al*., 2021; Li et al., 2022a; Martinez et al., 2023; Saunders et al., 2021; Walls et al., 2021) or N-terminal domain (NTD) (Joyce *et al*., 2021), as well as stabilized or scaffolded S2 stem immunogens (Hsieh et al., 2021; Kapingidza et al., 2023), have elicited broader antibody responses and demonstrated enhanced protection in animal challenge models. Other promising approaches are mosaic RBD nanoparticles combining RBDs from multiple CoVs (Cohen et al., 2021; Cohen et al., 2024; Walls *et al*., 2021; Zhang et al., 2022) and chimeric spike immunizations containing mixtures of RBDs, NTDs, or S2s from different CoVs (Chen et al., 2023a; Han et al., 2022; Martinez *et al*., 2021; Xu et al., 2022).

Another approach is to isolate broadly neutralizing antibodies from infected or vaccinated donors and use those to guide vaccine design - antibody-guided structure-based vaccine design (Kwong et al., 2020) or reverse vaccinology 2.0 (Burton, 2017). Since the emergence of SARS-CoV-2, numerous neutralizing monoclonal antibodies (nAbs) have been characterized and several bnAbs and specificities have been identified. These bnAbs show wide variation in the extent of their neutralization breadth from those that neutralize diverse SARS-CoV-2 VOCs, sarbecoviruses more generally, and even across β-coronaviruses (Brouwer et al., 2020; He et al., 2022a; Jennewein et al., 2021; Jette et al., 2021; Martinez et al., 2022; Pinto et al., 2020; Pinto et al., 2021a; Rogers et al., 2020; Tortorici et al., 2021; Zhou et al., 2023; Zhou et al., 2022). Three of the most promising sites to target for broad neutralization are the “silent face” of the receptor binding domain (RBD) (Song et al., 2025), the fusion peptide (Bianchini et al., 2023; Dacon et al., 2023; Low et al., 2022; Ng et al., 2022; Sun et al., 2022), and the stem-helix region of the spike S2 domain (Bianchini *et al*., 2023; Dacon *et al*., 2023; Hsieh *et al*., 2021; Hurlburt et al., 2022; Pinto *et al*., 2021a; Song et al., 2021; Wang et al., 2021a; Zhou *et al*., 2023; Zhou *et al*., 2022). Multiple human S2-specific mAbs with high cross-neutralizing potential have been isolated and characterized from infected donors (Dacon *et al*., 2023; Hurlburt *et al*., 2022; Pinto *et al*., 2021a; Song *et al*., 2021; Zhou *et al*., 2023). However, responses against these conserved bnAb target sites are infrequently elicited in natural infection or by current SARS-CoV-2 vaccines. Designed targeted vaccination strategies are thus needed to induce broadly protective bnAb responses.

In this study, we exploited the conservation of the spike S2 region among betacoronaviruses by utilizing heterologous sequential immunization to encourage B cell immunofocusing to common conserved S2 bnAb epitopes. We immunized rhesus macaques with adjuvanted SARS-CoV-2 S-protein followed by heterologous boosts with adjuvanted MERS-CoV S-protein and a β-coronavirus S-protein cocktail. Responses following the prime were overwhelmingly directed to the RBD as revealed by serology, electron microscopy-based polyclonal epitope mapping (EMPEM), and B cell profiling. Heterologous boosting resulted in re-focusing of responses to the common conserved S2 region, including the stem-helix bnAb site. However, S2-directed cross-reactive antibodies did not appear to contribute to serum cross-neutralizing responses. To further analyze responses, we employed an unbiased mAb isolation approach based on the magnitude of B cell clonal expansion across the cohort. This approach revealed major targeting of conserved S2 subunit B cell epitopes after heterologous boost but yielded mostly non-neutralizing, cross-reactive binding antibodies. Among the most clonally expanded lineages, we did, however, find two S2 stem-helix bnAbs that neutralized a broad spectrum of β-coronaviruses including MERS-CoV. These S2 stem-helix bnAbs were protective *in vivo* and structural studies revealed commonalities with prior human S2 stem-helix bnAbs. Therefore, this study demonstrates that vaccination can elicit bnAbs spanning sarbeco- to merbecoviruses, providing proof-of-principle for broadly protective β-coronavirus vaccines and establishing non-human primates as a relevant model for evaluating such strategies.

## Results

### Serum neutralizing antibody responses following heterologous CoV S-protein boosting

We previously showed that prime-boost immunizations in rhesus macaques with a recombinant SARS-CoV-2 S-protein along with a saponin (SMNP) adjuvant induced a robust serum bnAb response against SARS-related viruses (He et al., 2022b). We described the longitudinal bnAb responses up until week 16 and showed that they were primarily directed to the spike RBD. In the current study, we followed up with these animals and boosted at week 38 with SMNP-adjuvanted heterologous merbecovirus MERS-CoV S-protein to favor broader B cell recall responses to common shared spike epitopes, for example in the S2 subunit (Figure 1a). The MERS-CoV spike boost resulted in a strong MERS-CoV specific serum nAb response (Figure 1b) but nAb responses to SARS-CoV-2 on the other hand were not boosted. As previously described for the earlier time points (He *et al*., 2022b), we found no differences between the dose-escalated (DE) and bolus groups (Bolus) (Figures S1, S2, S3). The groups were combined for the remainder of this study. We then further boosted the animals at week 46 with a cocktail of three β-CoV S-proteins; MERS-CoV, HCoV-HKU1 and HCoV-OC43. HCoV-HKU1 and HCoV-OC43 are endemic human β-coronaviruses and like MERS-CoV share cross-reactive epitopes with SARS-CoV-2 spike in the S2 subunit (Ladner et al., 2021; Wang *et al*., 2021a). For the week 38 MERS-CoV boost, the three β-CoV S-protein cocktail at week 46 did not recall the SARS-CoV-2 nAb titers (Figure 1b). The nAb titers against MERS-CoV were further elevated following the cocktail boost, suggesting a potential recall response to the MERS-CoV antigen.

**Figure 1.**
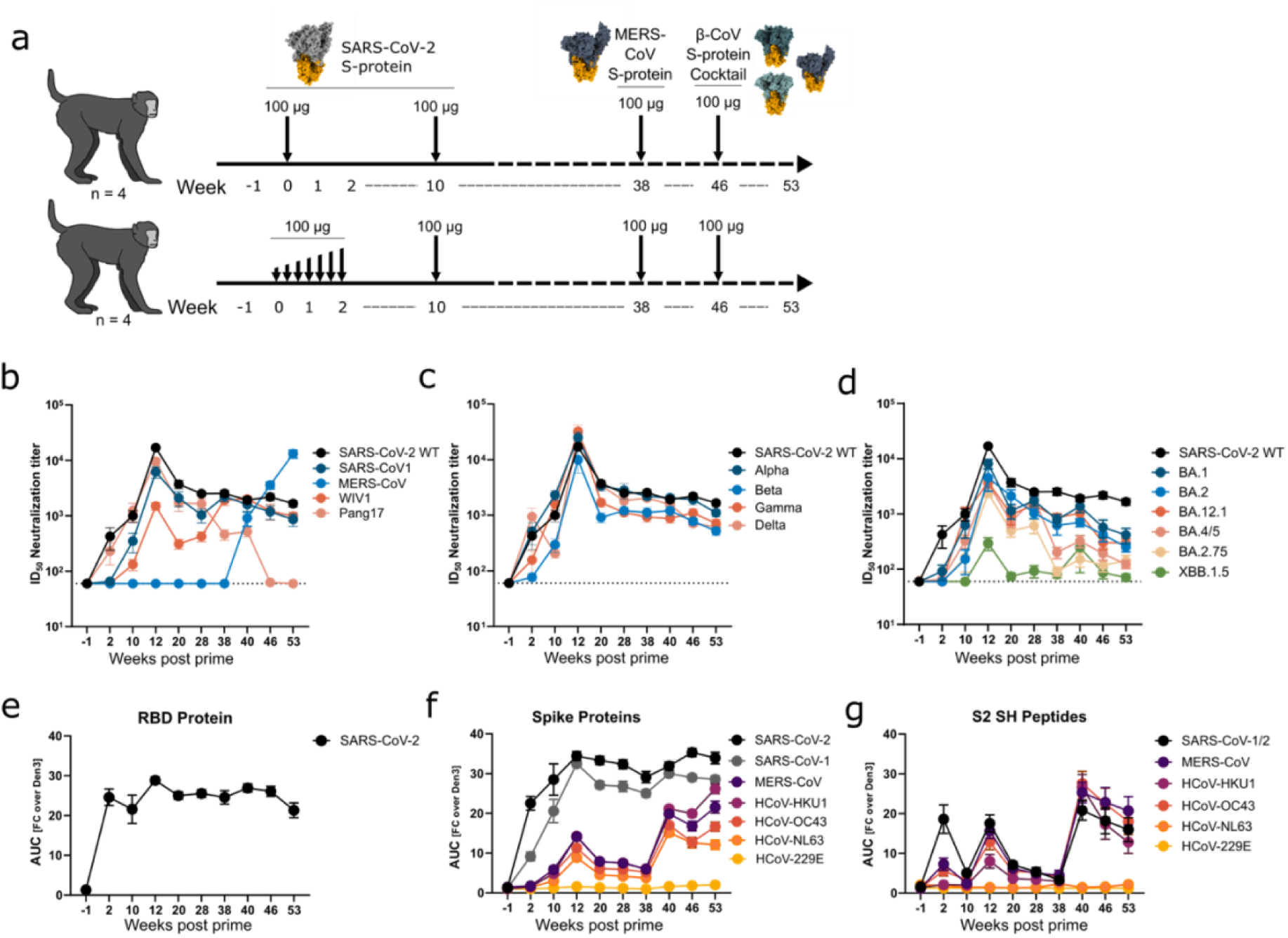
SARS-CoV-2 Spike-induced immune responses recalled by heterologous betacoronavirus spike boost elicit long-lasting humoral responses in rhesus macaques. **a)** Experimental design: rhesus macaques were primed with 100 μg SARS-CoV-2 spike protein in SMNP (saponin/MPLA nanoparticle) adjuvant at week 0. The priming immunization was given as either a single bolus dose (top, n=4) or as escalating dose (DE) (bottom, n=4) over 2 weeks. Both groups were boosted with 100 μg SARS-CoV-2 spike protein in SMNP at week 10, 100 μg MERS-CoV spike protein in SMNP at week 38, and a cocktail of 100 μg of the spike proteins of MERS-CoV, HCoV-HKU1 and HCoV-OC43 in SMNP at week 46. Structures depict spike proteins utilized for immunization. Serum and peripheral blood mononuclear cells (PBMCs) were sampled every 2 weeks from start to conclusion of the study at week 53. **b-d)** Serum ID_50_ neutralizing antibody titers were measured in S-protein–immunized rhesus macaques at select timepoints over the duration of the study. Serum neutralization was measured against pseudoviruses of select betacoronaviruses (b), SARS-CoV-2 variants of concern (VOCs) (c) and select SARS-CoV-2 Omicron variants (d). Data shown as a combination of both DE and bolus groups, with dots indicating mean and error bars representing standard error of the mean (SEM). Dashed line indicates the limit of detection. N=8 for all timepoints. **e-g)** NHP serum binding to receptor binding domains (e), spike proteins (f), and S2 stem helix peptides (g), of the indicated viruses was measured by ELISA at the indicated time points following priming immunization. Data are shown as area under curve (AUC) as fold change over negative control antibody Den3. Data are shown as a combination of both DE and bolus groups, with dots indicating mean and error bars representing SEM. N=8 for all time points.

Considering the breadth of nAb responses in more detail, the high titers of cross-neutralizing antibody responses against diverse sarbecoviruses (SARS-CoV-1, WIV1, and Pang17) were durable and persisted for more than six months (week 38) post the SARS-CoV-2 spike boost (week 10) (Figure 1b). We further tested the longitudinal immune sera for neutralization with SARS-CoV-2 early (Alpha, Beta, Gamma and Delta) and later Omicron (BA.1, BA.2, BA.12.1, BA.4/5, BA.2.75 and XBB.1.5) variants and observed that the early variants were neutralized as effectively as the WT SARS-CoV-2. Although high titers of nAbs against Omicron variants in the immune sera were observed for most variants, these titers dropped substantially over time. The XBB.1.5 Omicron variant, which is extensively mutated in the spike RBD region, was the most resistant virus against the immune sera at all time points. Overall, we observed strong and durable sarbecovirus spike RBD nAb responses induced by SARS-CoV-2 S-protein immunizations in rhesus macaques, but boosting with heterologous β-CoV S-proteins was largely ineffective at recalling these responses.

### Specificities of serum antibody responses following heterologous CoV S-protein boosting

To further understand the nature of the spike-induced antibody responses at prime and boost, we tested immune sera for binding to human coronavirus spike proteins and peptides. Consistent with the SARS-CoV-2 nAb responses above, we observed durable antibody binding titers to WT SARS-CoV-2 RBD (Figure 1c). These responses were induced by the first two SARS-CoV-2 S-protein immunizations and, consistent with SARS-CoV-2 specific nAbs, they remained unchanged upon heterologous β-CoV spike boosting. This latter result is likely due to the low sequence identity between the SARS-CoV-2 RBD and RBDs of other β-CoVs (72% for SARS-CoV-1; 24% for HKU-1; 23% for OC43; 19% for MERS-CoV) (Kaur et al., 2021)

Next, we tested the binding of immune sera to multiple α and β-CoV spikes (Figure 1c). As expected from the neutralization results above, we detected very strong and durable antibody binding responses to SARS-CoV-1 and SARS-CoV-2 S-proteins. We also detected moderate antibody binding to β-CoVs (MERS-CoV, HCoV-HKU1 and HCoV-OC43) post second immunization (week 10) with SARS-CoV-2 S-protein. These broader β-CoV responses were substantially elevated upon heterologous MERS-CoV (week 38) and three spike β-CoV spike cocktail boost (week 46) immunizations suggesting targeting of common conserved spike B cell epitopes. Further, weak but detectable antibody binding responses to phylogenetically distant α-CoV spike (HCoV-NL63) were observed particularly after the heterologous MERS β-CoV boosts suggesting possible targeting of common conserved B cell epitopes across α- and β-CoVs.

We further tested the binding of immune sera to S2 stem-helix peptides from α and β-CoVs. The S2 stem-helix region houses protective bnAb epitopes targeted by broad β-CoV bnAbs (Dacon et al., 2022; Pinto *et al*., 2021a; Zhou *et al*., 2023). Substantial binding to S2-stem helix peptides from β- but not α-coronavirus species (as expected due to the low sequence similarity between α- and β-CoVs) was observed especially after the heterologous MERS boost (Figure 1c).

Overall, the results suggested that heterologous coronavirus spike prime-boost immunizations in rhesus macaques increases serum B cell responses to multiple CoVs, but it is unclear to what extent this arises from priming of new responses and boosting of cross-binding responses.

### Epitope specificities of polyclonal serum antibody responses by EMPEM

To further investigate the epitopes targeted by the plasma anti-spike polyclonal antibodies (pAbs) at prime and boosts, we utilized electron microscopy-based polyclonal epitope mapping (EMPEM) (Bianchi et al., 2018). We complexed the antigen binding fragments (Fabs) of IgGs isolated from the immune plasma of 4 animals (K620, L603, DHIK and DHJB) with Wuhan SARS-CoV-2 spike to identify the epitope footprints of the elicited antibody responses at three time points; week 12, two weeks after the second immunization with SARS-CoV-2 S-protein; week 40, two weeks after the boost with MERS-CoV S-protein; and week 53, seven weeks after the final boost with the β-CoV S-proteins cocktail (Figure 1a, S4). EMPEM revealed that week 12 responses were dominated by RBD targeting antibodies (Figure 2a, b). RBD pAbs targeting epitopes around the receptor binding motif (RBM) were seen in all 4 NHPs, while pAbs targeting the RBD core (lower region of RBD) and the NTD were observed in 2 of the 4 animals (K620 and L603) tested (Figure 2a, b).

**Figure 2.**
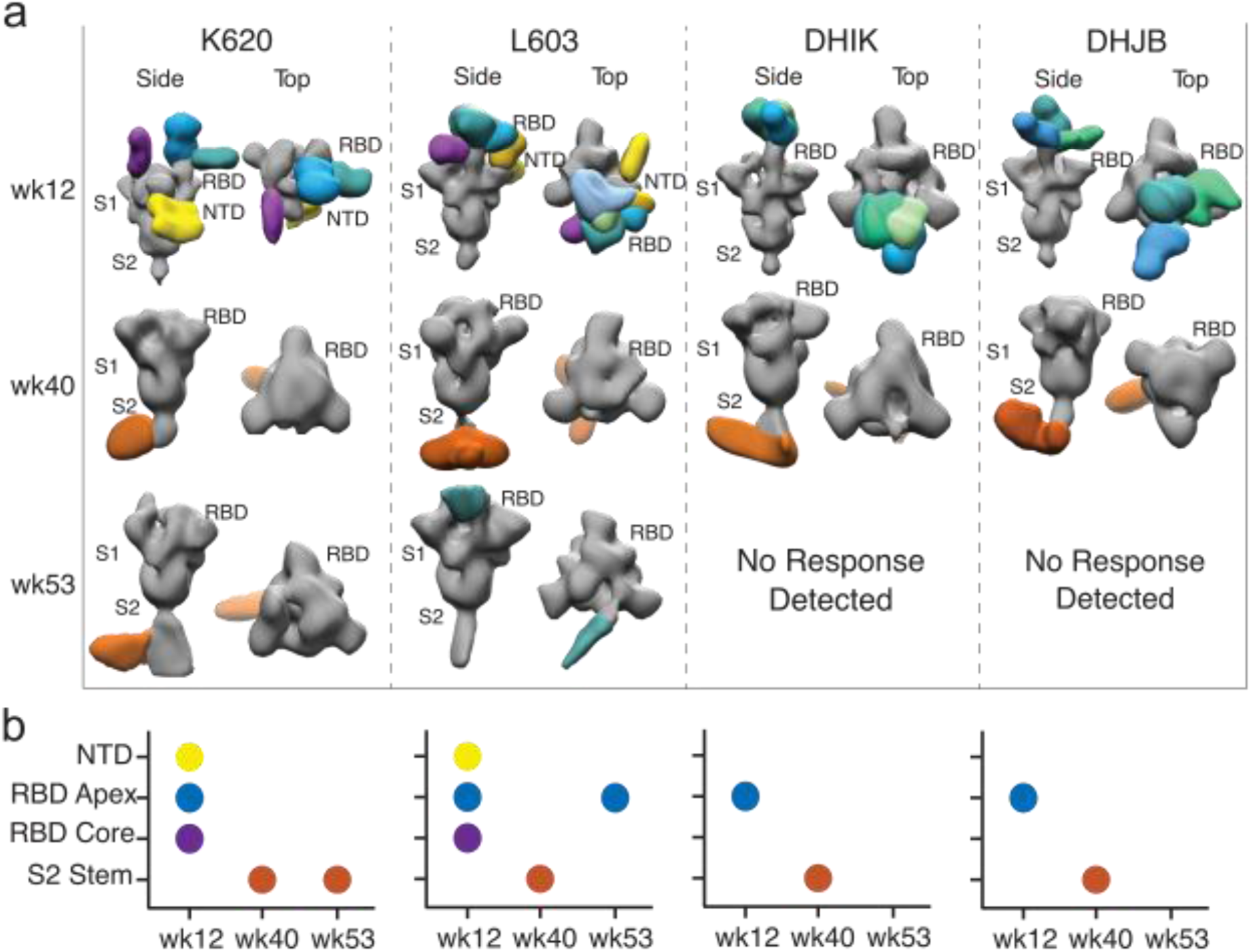
Analysis of serum antibody immune responses by EMPEM. **a)** Three-dimensional (3D) reconstructions generated from the negative stain electron microscopy (nsEM) images of polyclonal serum antibody Fabs of animals K620, L603, DHIK and DHJB isolated from serum taken at the indicated time points complexed with SARS-CoV-2 (Wuhan) spike protein. 2D class averages were used to generate 3D reconstructions. Colors of the Fabs indicate the following binding sites on the SARS-CoV-2 spike protein (grey): RBD apex (blue/green), RBD core (purple), NTD (yellow), and S2 stem helix (maroon). Individual Fabs are shown in shades of the base color. **b)** Summary plot of the data shown in a). Each plot represents the timepoints from a single animal as in a).

We next examined week 40 responses, which revealed S2 stem-helix pAbs as the major specificity elicited by the MERS-CoV spike boost (Figure 2a, b). We also observed varying levels of antibody-induced trimer dissociation in these datasets, which may reflect a shift from an immunodominant primary response towards more conserved epitopes that likely destabilize the spike trimer upon spike binding (Figure S5) (Song *et al*., 2025). To investigate this further, we processed all particles, including non-intact spikes, and performed masked 3D classification to eliminate alignment bias from the dominant S2 specificities. The resulting 3D reconstructions of partially dissociated spike trimers revealed Fab densities on the RBD (Figure S5). Although the low resolution of these reconstructions prevents conclusive identification of the RBD epitopes, the findings align with the presence of RBD-targeting antibodies at these time points. Following the heterologous spikes cocktail boost by week 53, EMPEM responses were more variable between animals with S2 targeting only observed clearly in one animal (K620), indicating that the spike cocktail boosting was likely inefficient at recalling B cell responses to the shared conserved epitopes.

Overall, EMPEM revealed targeting of the spike RBD epitopes with SARS-CoV-2 primary immunization, some immunofocusing to the common shared S2 epitopes after a heterologous spike boost, but evidence of more mixed responses on further boosting with a cocktail of spike immunogens.

### Coronavirus spike specific memory and plasma B cell responses

Next, we aimed to elucidate the B cell responses associated with each of the vaccination steps. Thus, we analyzed peripheral blood mononuclear cells (PBMCs) from three animals at weeks 10, 12, 40 and 53 (Figure 3, S6). We observed that the SARS-CoV-2 spike priming immunization induced robust SARS-CoV-2 spike protein and RBD targeting responses in both memory B cell (MBC) and plasma cell (PC, likely short lived plasmablast) compartments (Figure 3a, b). The response that developed after the initial priming until week 10 was successfully boosted by the week 10 homologous boost as shown by a significant increase in SARS-CoV-2 specific MBCs (Fig. 3b; p=0.002), and a limited increase in SARS-CoV-2 specific PCs. The heterologous boost at week 38 did not increase the numbers of SARS-CoV-2 spike-specific B cells and in fact the MBC numbers decreased markedly, although the numbers of PCs were maintained. By week 53, the numbers of both SARS-CoV-2 spike-specific MBCs and PCs in the blood had dropped considerably to levels below those seen at week 10 (Figure 3a, b). In accordance with our EMPEM and serum findings, these results indicate that the initial response in both B cell populations was directed against epitopes specific to SARS-CoV-2 and the heterologous boost spike proteins did not significantly boost cross-reactive B cell responses. The SARS-CoV-2 RBD protein-targeting MBC response stayed constant after the initial priming immunization as observed in the serum antibody responses, again pointing to failure to boost existing RBD-directed response by the heterologous boost (Figures 1, 2).

**Figure 3.**
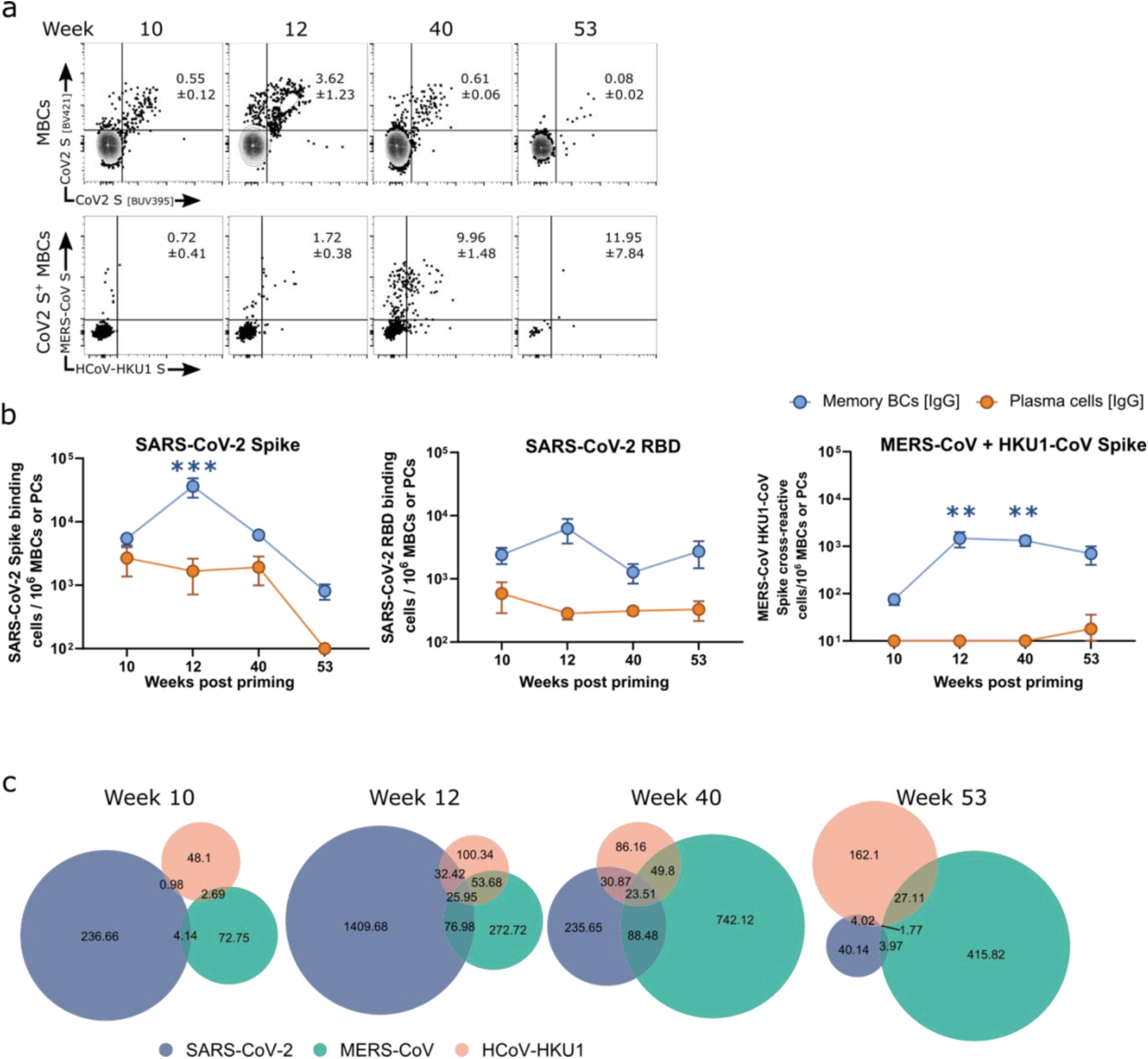
Antigen-specific B cell response to heterologous vaccination. Memory B cells are defined as CD8^-^ CD14^-^ CD4^-^ CD71^-^ CXCR5^-^ CD20^+^ CD38^+^ CD27^+^ IgG^+^; Plasma cells are defined as CD8^-^ CD14^-^ CD4^-^ CD20^-^ CD38high IgG^+^. **a)** Gating on SARS-CoV-2 spike-binding PBMC memory B cells (top) or MERS-CoV spike protein HCoV-HKU1 spike protein and SARS-CoV-2 spike protein cross-reactive PBMC memory B cells (bottom). Numbers shown are the mean percentage of cells in the indicated gates per parental population ± SEM. All graphs show equivalent cell numbers, except week 53; here all collected cells are shown. **b)** SARS-CoV-2 spike protein binding, SARS-CoV-2 RBD binding or MERS-CoV spike protein and HCoV-HKU1 spike protein cross-reactive memory B cells (blue) and plasma cells per 10^6^ total memory B cells or plasma cells in PBMCs. Data shown as mean ± SEM. Significance compared to week 10 was calculated by using two-way ANOVA with Holm-Šídák multiple comparison test. Significance is indicated by blue (PMBCs) or purple (PCs) asterisks above the data points. **c)** Venn diagram showing the number of PBMC memory B cells binding SARS-CoV-2 S protein, MERS-CoV S protein or HCoV-HKU1 S protein as determined by flow cytometry. Numbers shown are the indicated B cell populations per 10^6^ CD20^+^B cells. Size of circles is proportional to cell number within each graph. N=6 (N=3 dose escalation animals, N=3 bolus animals). *P < 0.05, **P < 0.01, ***P < 0.001, ****P < 0.0001, no asterisk: not significant (P > 0.05).

The priming immunization also induced low but detectable MERS-CoV/HCoV-HKU1 spike protein-directed responses (Figure 3b, c). While MERS-CoV/HCoV-HKU1-specificity was not accompanied by SARS-CoV-2 spike cross-reactivity at week 10, the boost at week 12 elicited cross-reactive MBCs and PCs (Figure 3b, c; MBCs: p=0.002) indicating targeting of common conserved spike sites between these viruses, such as the S2 region. At week 40, two weeks after the first heterologous boost, we observed a significant increase in MERS-CoV and HCoV-HKU.1 spike protein-specific MBCs (Figure 3b, c) (p=0.004), a robust fraction of which showed cross-reactivity to all three spike proteins (Figure 3c). Thus, not only was the heterologous boost capable of prompting the development of serum antibody responses and the appearance of short-lived S2-targeting PCs, it also selectively boosted S2-specific MBCs, suggesting the potential for long-term antibody protection.

At week 53, after the second heterologous boost, the SARS-CoV-2 directed response was dwarfed in comparison to the MERS-CoV spike/HCoV-HKU1 spike-specific responses in both MBCs and PCs (Figure 3b, c). The results show concordance between the effects of immunization on serum and memory responses.

### Binding, neutralization and immunogenetics of spike vaccine-elicited rhesus mAbs

At the completion of the vaccination protocol at week 53, we sacrificed the animals and obtained multiple tissues for further study. To generate mAbs, we isolated antigen-specific B cells from lymph node samples using SARS-CoV-2 S-protein as a bait (Figure 4a, S7). Briefly, we pooled lymph node samples from all eight vaccinated rhesus macaques, isolated SARS-CoV-2 S-protein binding IgG^+^ B cells through flow cytometric sorting, and processed the cells using the 10x Genomics Chromium platform. We sequenced the resulting libraries to analyze the heavy-light chain pairs of the antigen-sorted single B cells. We recovered 8435 antigen-specific B cells and identified the 50 largest expanded B cell lineages; reasoning that expanded lineages would correspond to interesting polyclonal Abs generated by vaccination. The most expanded B cell lineage, represented by CHM-1, possessed 661 clonally related members, and the least expanded lineage (CHM-50) had 12 members (Figure S8). MAbs representing 48 of the identified lineages were produced and characterized. 43 out of 48 mAbs showed strong binding by ELISA to soluble SARS-CoV-2 S-protein, which were used for the B cell enrichments, thus validating that our 10X Genomics B cells isolation method was ∼90% specific (Figure 4b, c). All mAbs that showed binding to SARS-CoV-2 exhibited cross-reactive binding to SARS-CoV-1 S-protein, except for four mAbs (Figure 4c). Approximately 65% of these mAbs displayed strong cross-reactive binding to S-proteins of other human β-HCoVs (MERS-CoV, HCoV-HKU1 and HCoV-OC43), but weak and much lower frequency binding to α-HCoV, HCoV-229E and rarely to HCoV-NL63 S-protein. A small fraction of mAbs showed binding to all seven human coronavirus S-proteins. Out of SARS-CoV-2 S-protein mAbs, six were specific to RBD as determined by binding to monomeric SARS-CoV-2 RBD protein. We also tested binding of all mAbs with 25mer S2 stem-helix peptides derived from seven human β-HCoV and α-HCoV spikes. Notably, two of the mAbs, CHM-16 and CHM-27 exhibited cross-reactive binding to β- but not α-HCoV S2 stem peptides. Overall, many mAbs showed binding to multiple β-HCoV spikes suggesting targeting of shared epitopes.

**Figure 4.**
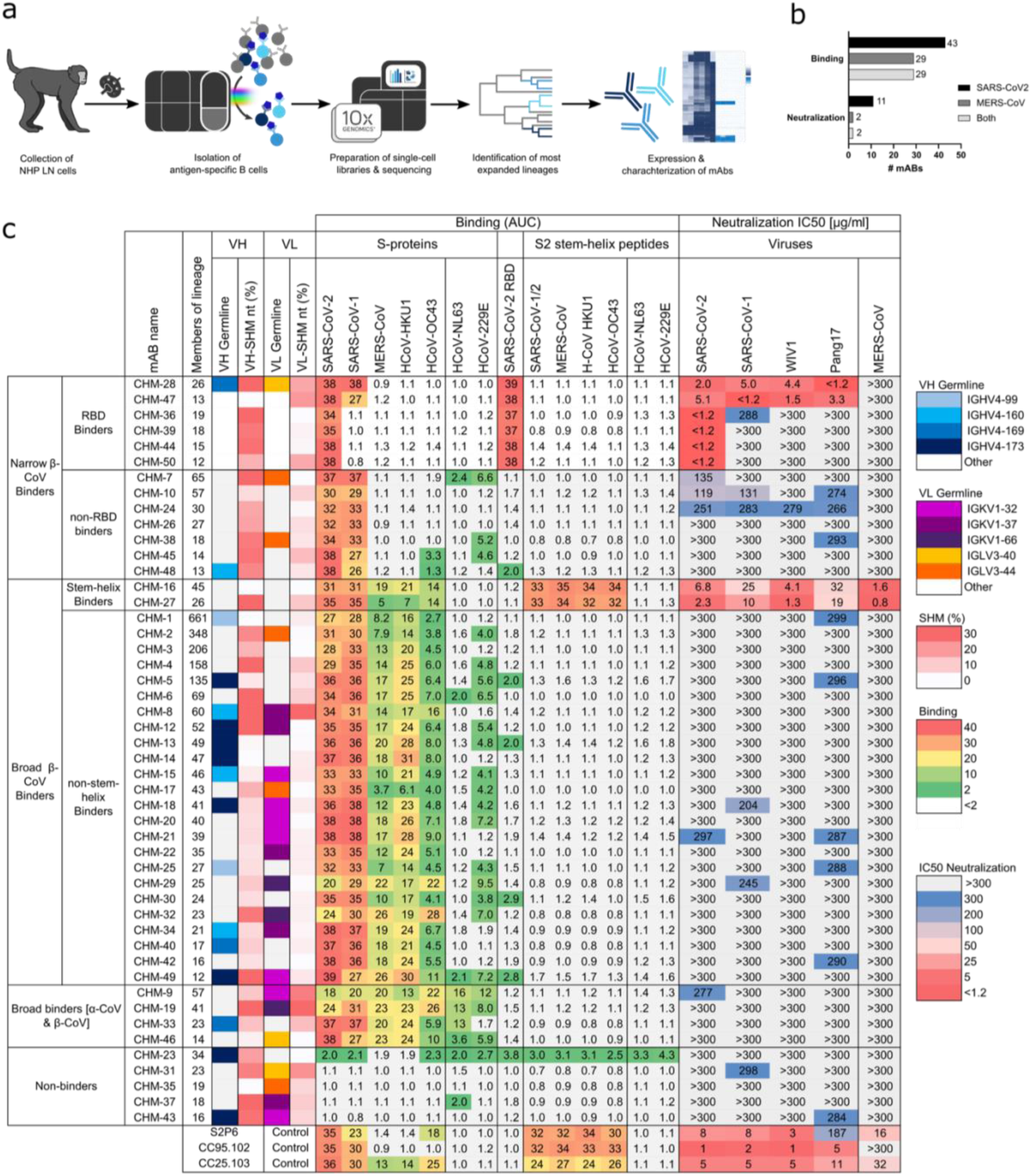
Expanded antibody lineages isolated from immunized NHPs allow analysis of the full spectrum of elicited antibodies. **a)** Experimental setup. Draining lymph node cells of all eight immunized NHPs were collected at week 53 and bulk sorted for IgG^+^ SARS-CoV-2 spike protein binding B cells. Single cell RNA-seq libraries were generated using 10X Genomics kits and subsequently sequenced. The resulting antibody sequences were analyzed for expanded lineages across animals. After identifying the 50 most highly expanded lineages, the most expanded clones per lineage were expressed as mAbs. 48 mAbs were expressed and characterized. Two lineages were not included in this study. **b)** Quantification of mAbs out of 48 total mAbs that either bind SARS-CoV-2 and/or MERS-CoV spike proteins or neutralize the corresponding pseudoviruses. **c)** A total of 48 mAbs were isolated using the method outlined above and grouped by binding and neutralization characteristics. The heatmap shows total members of antibody lineage as identified via single cell RNA-seq, IGHV germline gene usage [select expanded IGHV genes [shades of blue], other IGHV genes [gray]), IGLV germline gene usage (select expanded IGKV genes [shades of purple], select expanded IGLV genes [shades of orange], and other VL genes [gray]), and V-gene nucleotide somatic hypermutations (SHMs). ELISA binding of mAbs with β- and α-HCoV spike proteins and S2 stem-helix region peptides are shown as area under curve (AUC) as fold change over negative control mAb (Den3). Neutralization capabilities of mAbs against pseudoviruses of clade 1a (SARS-CoV-2 and Pang17), clade 1b (SARS-CoV-1, WIV1) sarbecoviruses, and MERS-CoV are shown as IC_50_ [μg/ml]. Controls for ELISA and neutralization assays are the previously described spike S2 stem-helix bnAbs CC40.8, S2P6, CC95.102, and CC25.103, as well as the negative control mAb Den3.

We next examined neutralization of mAbs against clade 1b (SARS-CoV-2 and Pang17) and clade 1a (SARS-CoV-1, and WIV1) ACE2-utilizing sarbecoviruses and MERS-CoV. All six of the spike RBD specific mAbs (CHM-28, CHM-47, CHM-36, CHM-39, CHM-44 and CHM-50) potently neutralized SARS-CoV-2 (geometric mean IC_50_: SARS-CoV-2 =0.8 µg/ml) (Figure 4c). Two of these RBD nAbs (CHM-28 and CHM-47) exhibited cross-neutralization with SARS-CoV-1, WIV1 and Pang17 sarbecoviruses but failed to neutralize MERS-CoV due to the RBD sequence divergence. Seven of the mAbs that showed strong binding to SARS-CoV-2 and SARS-CoV-1, but weak or no binding to β-CoV spikes (narrow binders), exhibited some very weak neutralization. Notably, two (CHM-16 and CHM-27) of the 26 “broad β-CoV binders’’ showed broad cross neutralization of sarbecoviruses and MERS-CoV and both bind to the S2 stem-helix bnAb site of the spike fusion machinery (Figure 4c). The IC_50_ neutralization potencies for these two bnAbs were moderate, typical of human S2 stem-helix bnAbs (Dacon *et al*., 2023; Zhou *et al*., 2023; Zhou *et al*., 2022). These two bnAbs represent, to our knowledge, the first examples of the sarbeco-/merbeco-virus S2 stem-helix bnAbs elicited by immunization in rhesus macaques.

### Macaque S2 stem bnAbs share epitope and immunogenetic features with human bnAbs

We then focused on the two bnAbs CHM-16 and CHM-27. We found neutralization across all sarbecoviruses (SARS-CoV-1, SARS-CoV-2, WIV1, and Pang17) and merbecoviruses (MERS-CoV) tested (Figure 5a). In addition, we observed consistently strong neutralization across key SARS-CoV-2 Omicron VOCs, demonstrating the broad cross-neutralization capabilities of the S2 stem-helix bnAbs. CHM-16 and CHM-27 both exhibited strong binding to SARS-CoV-1 and SARS-CoV-2 spike proteins by ELISA (Figure S9). MERS-CoV, HCoV-HKU1 and HCoV-OC43 spike proteins, however, were bound at moderate strength by CHM-16. CHM-27 showed binding to HCoV-OC43 only at high antibody concentrations (>2 µg/ml) and only weak binding to MERS-CoV and HCoV-HKU1. None of the bnAbs showed any binding to α-coronavirus spike proteins or SARS-CoV-2 RBD. When analyzing the binding abilities to S2 stem-helix (SH) peptides, CHM-16 and CHM-27 exhibited similar strong binding to the S2 SH peptides of SARS-CoV-1, SARS-CoV-2, MERS-CoV, and HCoV-OC43, suggesting S2 SH epitope targeting of these bnAbs (Figure S9). While CHM-16 showed similar high affinity for HCoV-HKU1 as for the aforementioned S2 SH peptides, CHM-27’s binding affinity for HCoV-HKU1 was comparatively reduced. The stronger binding of CHM-16 to the MERS-CoV and HCoV-HKU1 S2 SH peptide in comparison to spike proteins of the same viruses might be due to the limited epitope accessibility on the spike protein.

**Figure 5.**
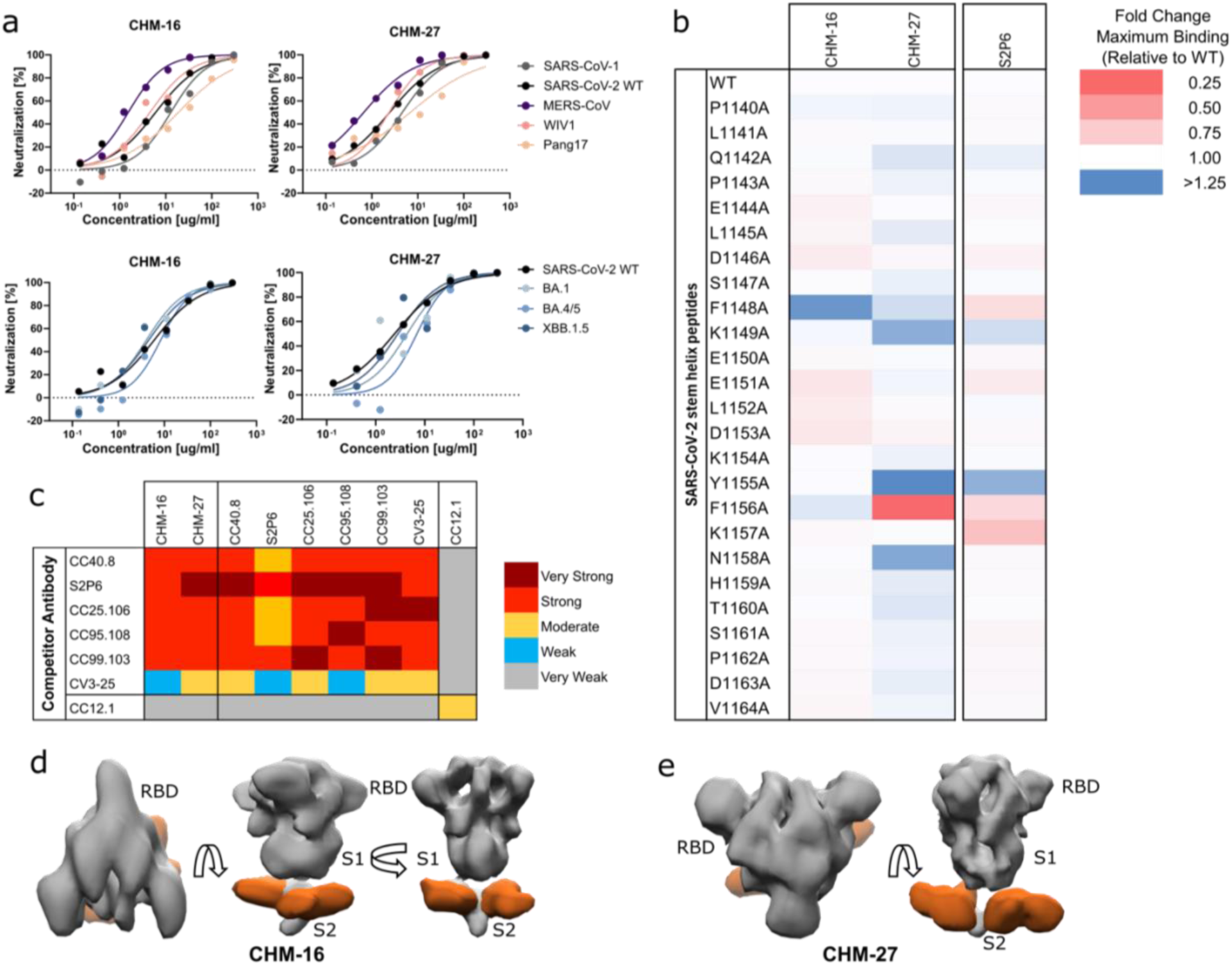
Characterization of isolated S2 spike stem-helix antibodies. **a)** Neutralization of pseudoviruses of the indicated β-coronaviruses (top) and SARS-CoV-2 variants (bottom) by CHM-16 and CHM-27. **b)** BLI-based epitope mapping of S2 stem-helix binding mAbs CHM-16 and CHM-27 using 25-mer alanine scan peptides covering the SARS-CoV-2 stem-helix region. Heatmap shows fold-changes in maximum BLI binding responses of mAb binding to SARS-CoV-2 stem-helix peptide alanine mutants, compared with the WT peptide. **c)** BLI-based competition epitope binning to compare binding sites on SARS-CoV-2 spike protein of CHM-16 and CHM-27 to known binding sites of S2 stem-helix bnAbs CC40.8, S2P6, CC25.106, CC95.108, CC99.103, and CV3-25. CC12.1 served as negative control. Binding levels (“Very strong” to “very weak”) are indicated by color. **d-e)** EM-based epitope mapping 3D reconstruction of monoclonal CHM-16 (d) and CHM-27 (e) Fabs (orange) complexed with SARS-CoV-2 spike protein in multiple orientations. 3D reconstruction was generated from 2D class averages.

To further determine the specific epitopes of CHM-16 and CHM-27, we generated alanine scanning mutants of the SARS-CoV-2 stem peptide and performed binding assays (Figure 5b). We found dependence on D1146, E1151, L1152, and D1153 for CHM-16 and stronger dependence of CHM-27 on F1156, indicating slightly differing epitope specificities for CHM-16 and CHM-27 in accordance with earlier studies on human S2 stem bnAbs (Dacon *et al*., 2022; Pinto *et al*., 2021a; Zhou *et al*., 2023). For additional epitope mapping of CHM-16 and CHM-27, we utilized epitope binning via bio-layer interferometry (BLI) with well-characterized S2 stem-helix bnAbs, CC40.8 (Zhou *et al*., 2022), S2P6 (Pinto et al., 2021b), CC25.106, CC95.108, CC99.103 (Zhou *et al*., 2023), and CV3-25 (Li et al., 2022b). With exception of CV3-25, which competed moderately, all bnAbs competed strongly or very strongly with CHM-16 and CHM-27 (Figure 5c), suggesting that CHM-16 and CHM-27 share overlapping epitopes with the panel of competing human S2 stem bnAbs. Based on these results and as described recently for human S2 stem-helix bnAbs (Zhou *et al*., 2023), CHM-16 and CHM-27 target a highly conserved region containing the hydrophobic core residues of the spike fusion machinery. Single-particle negative-stain electron microscopy confirmed that both CHM-16 and CHM-27 target the S2 stem-helix region of the SARS-CoV-2 S-protein (Figures 5d, e, S10).

Collectively, the results suggest that rhesus S2 stem-helix bnAbs target similar epitope elements in the stem-helix fusion machinery and use immunogenic features that closely resemble humans. Thus, rhesus macaques could serve as a favored animal model to evaluate S2 stem-based vaccines.

### Structural studies of macaque S2 stem-helix broadly neutralizing antibodies

To elucidate potential mechanisms of neutralization of CHM-16 and CHM-27 in molecular detail, we determined crystal structures of bnAb CHM-16 in complex with the SARS-CoV-2 S2 stem peptide and bnAb CHM-27 in complex with the MERS-CoV S2 stem peptide, at resolutions of 3.07 Å and 1.80 Å, respectively (Figure 6 and Table S1). The S2 stem peptides in both structures adopt a largely helical conformation and form extensive interactions with the antibodies. For CHM-16, CDRs H1, H2, H3, L1, and L3 interact with the SARS-CoV-2 S2 stem peptide. For CHM-27, all six CDR loops as well as light-chain framework 3 (FRL3) interact with the MERS-CoV S2 stem peptide (Figure 6a, b (far left panel)). For both antibodies, the heavy and light chains form a hydrophobic groove to accommodate the S2 stem helix (Figure 6a, b (middle left panel)), where 857 Å^2^ and 966 Å^2^ of surface is buried by CHM-16 and CHM-27, respectively, to cover approximately 44% and 55% of the total peptide surface area (Figure 6a, b (far right panel)). Superimposition of the CHM-16/S2 and CHM-27/S2 stem Fab structures onto SARS-CoV-2 spike proteins in both pre- and post-fusion states revealed steric clashes in both conformations (Figure S11). This suggests a potential neutralization mechanism involving disruption of the S2 stem three-helix bundle, with antibody binding likely occurring at an intermediate state or a locally rearranged conformation of the spike, as previously proposed for human S2 stem-helix bnAbs. (Pinto *et al*., 2021a; Zhou *et al*., 2023; Zhou *et al*., 2022).

**Figure 6.**
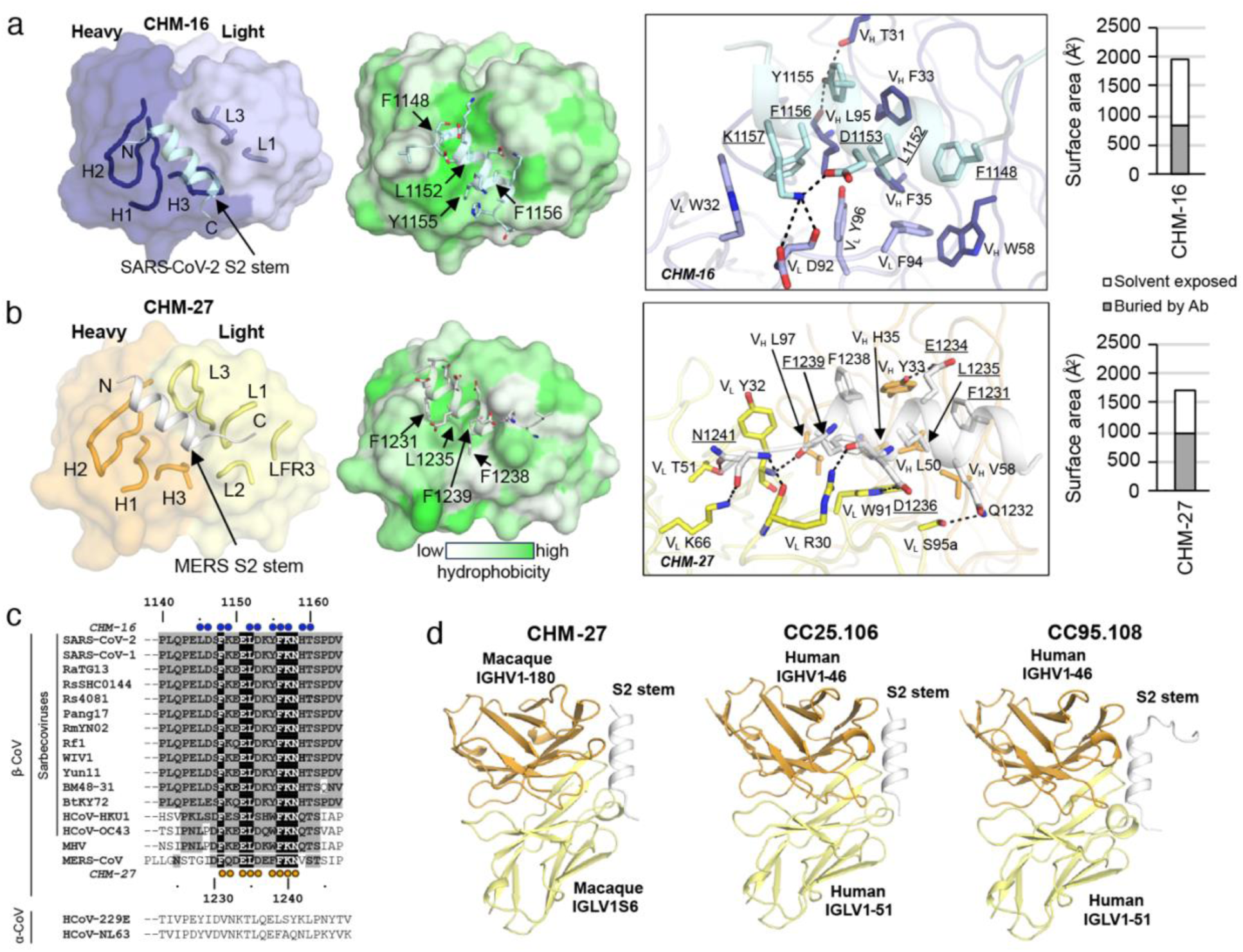
Crystal structures of mAbs CHM-16 and CHM-27 in complex with S2 stem peptides and comparison with other stem-helix mAbs. Heavy and light chains of CHM-16 are shown in blue and lavender, respectively, and the SARS-CoV-2 stem peptide is in cyan. Heavy and light chains of CHM-27 are shown in orange and yellow, respectively, and the MERS-CoV stem peptide in white. **a)** CHM-16 in complex with SARS-CoV-2 S2 stem peptide. Far left: overall view of the CHM-16-peptide structure at 3.07-Å resolution. Middle left: The stem-helix peptides insert into the hydrophobic grooves formed by the heavy and light chains of CHM-16. Surfaces of CHM-16 are color-coded by hydrophobicity [calculated by Color h (https://pymolwiki.org/index.php/Color_h)]. Middle right: detailed atomic interactions between the S2 stem peptides and antibody CHM-16. Kabat numbering is applied to the antibody. Numbering of the peptides is based on SARS-CoV-2 and MERS-CoV numbering. Hydrogen bonds and salt bridges are represented by black dashed lines. Conserved identical residues between SARS-CoV-2 and MERS-CoV are underlined. Far right: surface area of S2 stem peptide buried by CHM-16. Solvent exposed and buried areas were calculated with Proteins, Interfaces, Structures and Assemblies (PISA) (Krissinel and Henrick, 2007). **b)** CHM-27 in complex with MERS-CoV S2 stem peptide. Far left: overall view of the CHM-27-peptide structure at 1.8-Å resolution. Middle left: the stem-helix peptides insert into the hydrophobic grooves formed by the heavy and light chains of CHM-16. Surfaces of CHM-27 are color-coded by hydrophobicity [calculated by Color h (https://pymolwiki.org/index.php/Color_h)]. Middle right: detailed atomic interactions between the S2 stem peptides and antibody CHM-27. Kabat numbering is applied to the antibody. Numbering of the peptides is based on SARS-CoV-2 and MERS-CoV numbering. Hydrogen bonds and salt bridges are represented by black dashed lines. Conserved identical residues between SARS-CoV-2 and MERS-CoV are underlined. Far right: surface area of S2 stem peptide buried by CHM-27. Solvent exposed and buried areas were calculated with Proteins, Interfaces, Structures and Assemblies (PISA) (Krissinel and Henrick, 2007). **c)** Sequence alignment of the S2 stem region of β- and a-coronaviruses. Conserved identical residues in β-coronaviruses are highlighted with black boxes, and similar residues are in grey boxes [amino acids that scored greater than or equal to 0 in the BLOSUM62 alignment score matrix(Henikoff and Henikoff, 1992) were counted as similar here]. Residues of the SARS-CoV-2 S2 stem region that interact with CHM-16 are highlighted by blue circles above the alignment. Residues of the MERS-CoV S2 stem region interacting with CHM-27 are highlighted by orange circles below the alignment. Interaction is defined as distance between any atom in an amino acid of the peptide and the antibody of less than 4 Å. **d)** Structures of macaque mAb CHM-27 in complex with MERS-CoV S2 stem helix, human mAb CC25.106 in complex with SARS-CoV-2 S2 stem helix (PDB 8DGU), and mAb CC95.108 in complex with HCoV-HKU1 S2 stem helix (PDB 8DGW). All of the bound peptides are in the same orientation. Putative germline genes of the antibodies are labeled next to each chain. Heavy and light chains of antibodies are shown in orange and yellow, respectively. The stem-helix peptides are shown as white cartoons. For clarity, only variable domains of Fabs are shown.

The S2 stem-helix is highly conserved across β-coronaviruses (Figure 6c) and conserved residues contribute extensively to the recognition by both CHM-16 and CHM-27 (Figure 6a, b (middle right panel), explaining their broad reactivity to a variety of β-coronaviruses (Figure 4). For CHM-16, paratopic residues V_H_ F33, F35, W58, L95, and V_L_ W32, F94, Y96 form a groove to accommodate the hydrophobic face of the SARS-CoV-2 S2 stem helix that is comprised of aromatic and aliphatic residues F1148, L1152, Y1155 and F1156 (SARS-CoV-2 numbering). V_H_ T31 and L95 form hydrogen bonds (HB) with Y1155. V_L_ D92 engages K1157 through a HB and a salt bridge. K1157 also makes a salt bridge with D1153. For CHM-27, paratopic residues V_H_ Y33, L50, V58, L97, and V_L_ W91 form a groove to accommodate the hydrophobic face of the MERS-CoV S2 stem-helix that is comprised of conserved aromatic and aliphatic residues F1231, L1235, F1238, and F1239 (MERS-CoV numbering). V_L_ S95a, V_H_ Y33, V_L_ W91, and Y32 main-chain hydrogen bond with Q1232, E1234, D1236, and F1239 main-chain, respectively. V_L_ R30, T51, and K66 hydrogen bond with the main chain and side chain of N1241.

We next compared the crystal structures of the macaque antibodies CHM-16 and CHM-27 with previously published human anti-S2 stem antibodies (Figure S12). Strikingly, antibody CHM-27 elicited from an immunized naïve macaque targets the S2 stem helix in a nearly identical angle of approach as human IGHV1-46/IGLV1-51 public antibodies CC25.106 and CC95.108 (Zhou *et al*., 2023) (Figure 6d and Fig. S12). We then compared the nucleotide sequences of CHM-27 (encoded by macaque IGV genes IGHV1-180 and IGLV1S6) with human IGV germline genes by searching the IMGT database (Brochet et al., 2008). The most similar human IGHV germline genes to CHM-27 are indeed the ones that encode human public antibodies, IGHV1-46 and IGLV1-51. We then aligned the sequences of the macaque antibody CHM-27 and human antibodies CC25.106 and CC95.108 with their respective germline gene sequences (Figure S13a, b), and compared the detailed interactions of CHM-27, CC25.106 and CC95.108 with their respective peptide antigens (MERS-CoV, SARS-CoV-2, and HCoV-HKU1) (Figure S14). The structures showed that many of the paratopic residues involved in recognizing the S2 stem helices are conserved and are germline-encoded among all of these stem-specific antibodies from macaque and human. For example, V_H_ Y33, H35, and V_L_ W91 that interact with the S2 stem helices are encoded by the macaque and human germline genes. V_H_ W47 stabilizes V_H_ H35 and V_L_ W91, which interact with a Phe (F1239, F1156, and F1242). Conserved germline-encoded residues V_L_ Y/F32 and K66 form important interactions with an Asn (N1241, N1158, and N1244) at the C-term of the epitope. Macaque IGHV1-180 also encodes a germline residue V_H_ L50, similar to the human IGHV1-46 germline residue V_H_ I50, which are involved in interaction with the S2 stem helix. The human IGHV1-46 antibodies elicited a convergent somatic hypermutation V_H_ S56N, which corresponds to the macaque germline V_H_ N56, and forms a π-π interactions with a Phe (F1231, F1148, and F1234) (Figure S14 and S13a, b).

### Macaque S2 stem bnAbs show robust protection against SARS-CoV-2 challenge

To determine the protective efficacy of CHM-16 and CHM-27, we intraperitoneally (i.p.) treated groups (8-11 animal per group) of K18 mice expressing human ACE2 receptors with antibodies at 300µg per animal followed by a SARS-CoV-2 (USA-WA1/2020) challenge (Figure 7a). The SARS-CoV-2 challenge virus was intranasally (i.n.) administered 12 hours post-antibody infusion at a virus dose of 2 × 10^4^ focus forming units (FFU) (Figure 7a). Spike RBD nAb, CC12.1 (positive control), or a Zika-specific antibody, SMZAb1 (negative control) were administered into the control animal groups. The animals were weighed daily to monitor weight changes, which is an indicator of disease progression due to infection. All animals were euthanized on day 6 and lung tissues were collected to determine the virus titers by two methods: i) focus forming unit assays and, ii) quantitative polymerase chain reaction (qPCR). The S2 stem-helix bnAb treated animals showed significantly reduced weight loss as compared to the negative control antibody SMZAb1-treated animals at day 6 (P<0.0001, Figure 7b, c), confirming the protective role of S2 stem-helix bnAbs against SARS-CoV-2. Both S2 stem-helix bnAbs showed comparably significant protective effects from day 4 until the end of the study. The positive control RBD nAb, CC12.1 also significantly protected animals against weight loss (P<0.0001, Figure 7b, c). Infectious viral titers and viral RNA copies in day 6 lung tissues were significantly reduced in S2 antibody-treated animals compared to the SMZAb1 control group animals (viral titers: P<0.0001; viral load: P= 0.0164; Figure 7d, e). Although, we did not here evaluate protection against multiple β-coronaviruses, previous studies have shown that S2 nAb titers are predictive of protection (Zhou *et al*., 2023). As similar nAb IC50s were observed for CHM-16 and CHM-27 against a range of β-CoVs, we anticipated that protection will be provided equivalently against these viruses.

**Figure 7.**
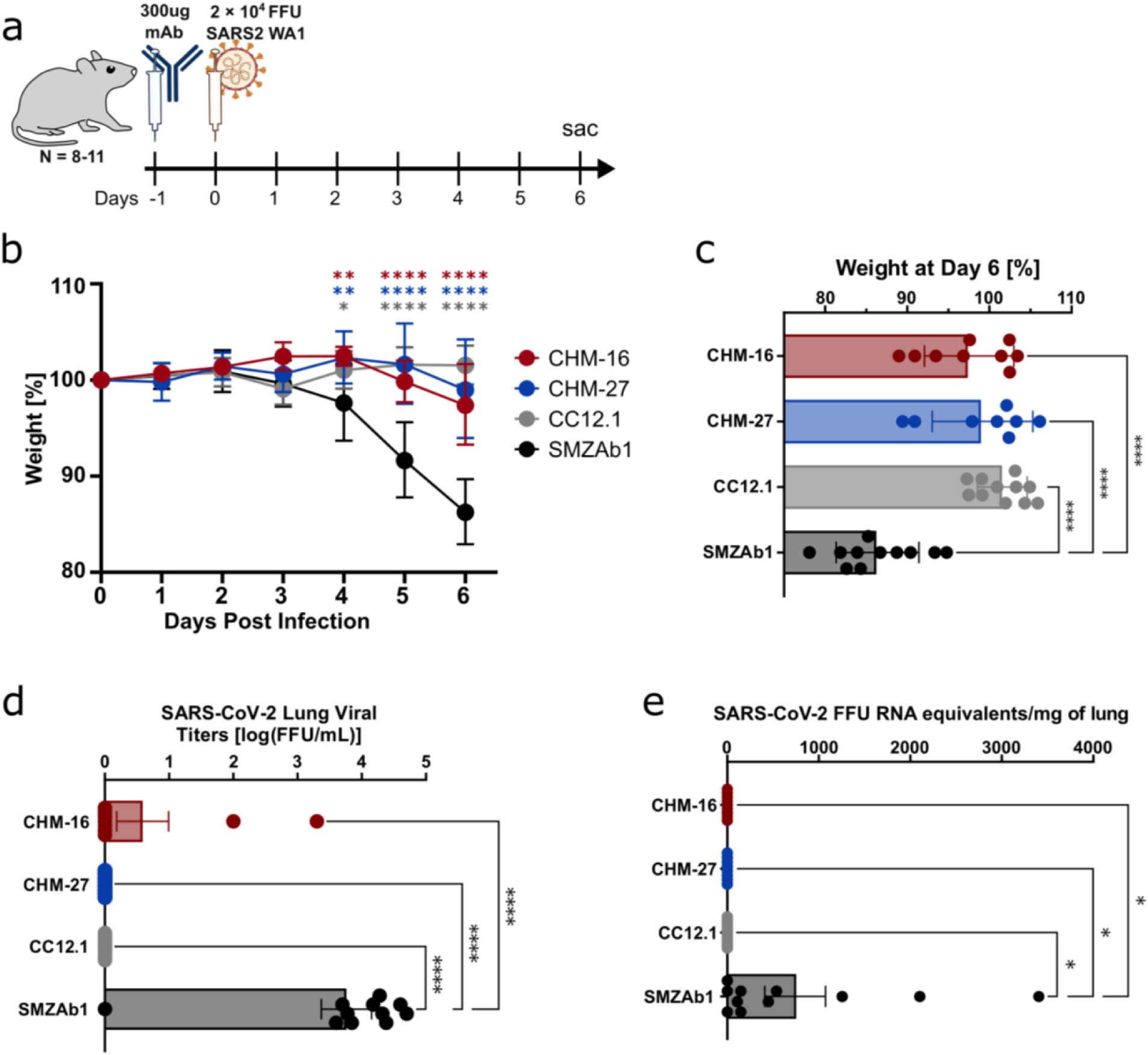
S2 spike stem-helix antibodies protect against SARS-CoV-2 in murine challenge model. **a)** K18 mice e xpressing human ACE2 receptors were administered 300ug of mAbs CHM-16, CHM-27 or positive (CC12.1) or negative (SMZAb1) control mAbs at day -1 i.p. 12h after mAb infusion, mice were challenged with SARS-CoV-2 virus (USA-WA1/2020) at day 0. Mice were weighed daily and the study was concluded on day 6 post-challenge. **b)** Geometric mean of percent change of weight compared to baseline for each group of mice. Significance of weight loss compared to negative control (SMZAb1) was calculated by using two-way ANOVA with Holm-Šídák multiple comparison test. Asterisks indicating significance are shown above the data points and are color coded by group. **c)** Shown is mean ± SEM of weight change at day 6 in percent of weight at day 0 of each individual mouse. Significance compared to negative control (SMZAb1) was calculated by using one-way ANOVA with Holm-Šídák multiple comparison test. **d)** Lung virus titers of mice at day 6 measured as focus forming units (FFU). Significance was determined by using one-way ANOVA with Holm-Šídák multiple comparison test compared to the negative control (SMZAb1). **e)** Viral RNA found in lungs of mice at day 6 post challenge as measured via qPCR. Significance calculation was performed by using one-way ANOVA with Holm-Šídák multiple comparison test compared to the negative control (SMZAb1).**a)** CHM-16: n=9; CHM-27: n=8; n=11 for CC12.1 and SMZAb1. *P < 0.05, **P < 0.01, ***P < 0.001, ****P < 0.0001, ns: not significant (P > 0.05).

## Discussion

Targeting conserved spike bnAb sites holds promise for developing broad vaccine strategies capable of addressing current SARS-CoV-2 variants as well as potential future pandemic coronaviruses (Chen et al., 2023b). While three highly conserved bnAb sites on the SARS-CoV-2 spike have been previously identified (Bianchini *et al*., 2023; Dacon *et al*., 2023; Hsieh *et al*., 2021; Pinto *et al*., 2021a; Song *et al*., 2021; Song *et al*., 2025; Wang *et al*., 2021a; Zhou *et al*., 2023; Zhou *et al*., 2022), these epitopes are partially occluded on the prefusion trimeric spike, limiting the frequency with which bnAbs targeting them are elicited through natural infection or repeated immunization with a single spike trimer immunogen. This highlights the need for rationally designed immunogens and immunization strategies to effectively induce such broadly neutralizing responses.

In this study, we focused on targeting the conserved S2 subunit on betacoronaviruses and immunized rhesus macaques with adjuvanted SARS-CoV-2 S protein, followed by heterologous boosts with MERS-CoV S protein and a β-coronavirus spike cocktail to favor B cell responses to common conserved epitopes. This approach induced serum antibodies with cross-reactive binding to divergent β-coronavirus spikes, although serum-level broad betacoronavirus-neutralization was largely absent. Importantly, using an unbiased strategy to isolate monoclonal antibodies based on major B cell clonal expansions, we identified two S2 stem-helix antibodies that were broadly neutralizing across sarbeco- and merbeco-coronaviruses, and conferred protection against SARS-CoV-2 challenge in an *in vivo* mouse model. These antibodies also share structural and sequence features with human S2 stem-helix bnAbs, including conserved paratopic residues, and one macaque antibody (CHM-27) adopts a nearly identical angle of approach to the human IGHV1-46/IGLV1-51 public class (Zhou *et al*., 2023), despite only 81.4% germline nucleotide identity.

To our knowledge, this is the first demonstration of S2 stem-helix bnAbs elicited by vaccination in rhesus macaques, highlighting both the translational relevance of NHP models and the potential to recapitulate human bnAb responses through rational immunization. These results demonstrate proof-of-principle for sequential heterologous spike immunization to engage B cell responses targeting common conserved, cross-reactive S2 epitopes. However, the majority of S2-directed responses remain non-neutralizing due to the cryptic nature of the stem-helix epitope on the prefusion spike. Our findings highlights that a major barrier to eliciting S2-stem directed bnAbs is the cryptic nature of the stem-helix epitope within the pre-fusion spike, where limited accessibility constrains effective B cell activation. To overcome this barrier, rational vaccine design strategies that focus immune responses on the bnAb epitope are required. Epitope-based immunogens displayed on engineered scaffolds (Correia et al., 2014; Kapingidza *et al*., 2023) or stabilized stem immunogens (Hsieh *et al*., 2021) or multivalent nanoparticles (Kim et al., 2022) can present the S2 stem-helix site in an optimized configuration, thereby increasing the likelihood of priming bnAb precursor B cells. Once primed, these responses could be further shaped through sequential boosting with native-like spike immunogens that transiently or partially expose the S2 bnAb site, effectively training memory B cells to recognize this epitope in its native spike configuration. Such a stepwise immunization strategy has the potential to drive affinity maturation, enhance breadth, and ultimately elicit broadly protective S2 stem-helix bnAb responses against diverse betacoronaviruses.

Overall, our study demonstrates that heterologous coronavirus spike immunizations can elicit structurally and functionally relevant cross-clade β-coronavirus S2 stem bnAbs in non-human primates and provides a potential strategy for overcoming the limitations of cryptic, conserved epitopes in eliciting broadly protective coronavirus immunity.

## Acknowledgements

This work was supported by National Institutes of Health-(NIH), National Institute of Allergy and Infectious Diseases-(NIAID) awards, R01AI170928 (R.A.) and CHAVD UM1 AI44462 (A.B.W., D.R.B.) and the Bill and Melinda Gates Foundation INV-004923 (A.B.W., I.A.W., D.R.B.). We thank Henry Tien for technical support with the crystallization robot. We are grateful to the staff of the National Synchrotron Light Source II (NSLS-II) beamlines 17-ID-1 and 17-ID-2. NSLS-II is a U.S. Department of Energy Office of Science User Facility. This research used resources of the National Synchrotron Light Source II, a U.S. Department of Energy (DOE) Office of Science User Facility operated for the DOE Office of Science by Brookhaven National Laboratory under Contract No. DE-SC0012704.

## Author contributions

K.D., T.C., Z.F., R.N.L., S.B., M.Y., A.B.W., B.B., I.A.W., D.R.B., and R.A. conceived and designed the study. K.D., W.-t.H., S.Callaghan, G.S., W.R., and M.F. conducted animal immunization studies and processed plasma samples. K.K.M., E.B.A., M.S., and M. Melo prepared the SMNP adjuvant. K.D., T.C., W.-t.H., S. Callaghan, and N.M. performed BLI, ELISA, virus preparation, neutralization, isolation, and characterization of animal sera and the resulting monoclonal antibodies. T.C., G.S., and W.-t.H. prepared virus mutant plasmids. J.H., R.M., P.S., Y.S., and B.B. performed the immunogenetic analysis of the antibodies. K.D., T.C., G.A., L.V., X.L., P.Y., F.A., M.Makhdoomi., and E.W.-G. prepared the monoclonal antibody, spike, and RBD proteins. R.N.L., S.B., and J.L.T. conducted negative stain electron microscopy studies. Z.F. and M.Y. determined crystal structure of the antibody-antigen complexes. N.B., E.G. and T.R. performed the murine challenge study. K.D., T.C., Z.F., R.N.L., J.H., S.B., M.Y., N.B., E.G., W.-t.H., S. Callaghan, G.A., L.V., X.L., J.L.T., R.M., G.S., N.M., P.S., P.Y., F.A., M. Makhdoomi, Y.S., S. Crotty, D.J.I., T.R., A.B.W., B.B., I.A.W., D.R.B., and R.A. designed the experiments and/or analyzed the data. K.D., T.C., Z.F., R.N.L., S.B., M.Y., A.B.W, B.B., I.A.W., D.R.B., and R.A. wrote the paper, and all authors reviewed and edited the paper.

## STAR+METHODS

## EXPERIMENTAL MODEL AND STUDY PARTICIPANT DETAILS

### Cell lines

FreeStyle 293-F cells (Thermo Fisher Scientific Cat# R79007) were maintained in FreeStyle 293 Expression Medium (Gibco Cat# 12338018). Expi293F cells (Gibco Cat# A14527) were maintained in Expi293 Expression Medium (Gibco Cat# A1435101). Suspension FreeStyle 293-F cells and Expi293F cells were incubated in the shaker at 150 rpm, 37°C, 8% CO_2_. High Five cells (Thermo Fisher Scientific Cat# B85502) at an MOI of 5 to 10. Infected High Five cells were incubated at 28 °C in Insect-XPRESS protein-free insect cell medium (Lonza Bioscience Cat# 12-730Q) with shaking at 110 rpm for 72 h for protein expression. Adherent HEK293Tcells and HeLa-ACE2 cells were grown in Dulbecco’s Modified Eagle Medium (DMEM) with 10% heat-inactivated FBS, 4mM L-Glutamine and 1% Penicillin-Streptomycin, maintaining in the incubator at 37°C, 5% CO_2_. The stable hACE2-expressing HeLa cell line was developed by transducing hACE2 into HeLa cells (ATCC CCL-2) using a lentivirus system. Cells with stable and high hACE2 expression were selected to be used for the pseudovirus neutralization assay. The cells were authenticated using STR profiling and are routinely tested and are free of any mycoplasma contamination.

## METHOD DETAILS

### Ethics statement

The rhesus macaque animal studies were conducted at Alphagenesis after protocol approval by Alphagenesis Institutional Animal Care and Use (IACUC) under approval number AUP 19-10. The murine animal studies were conducted at The Scripps Research Institute after IACUC approval under approval number 20-0003. At both facilities, animals were kept, immunized, and bled in compliance with guidelines outlined by the Animal Welfare Act and Guide for the Care and Use of Laboratory Animals (National Research Council, 1996).

### Immunization and sampling

The immunization regimen for the first ten weeks of this trial, along with the detailed preparation of the SMNP adjuvant, was described previously (He *et al*., 2022b) Briefly, eight healthy outbred rhesus macaques (Macaca mulatta) of Indian origin, aged 3-5 years and divided evenly in gender, were separated into a bolus prime group (group 1) and a dose escalation group (group 2). For every immunization in both groups, the doses were administered subcutaneously, with each dose being split in half and administered bilaterally to the left and right mid-thigh. The bolus prime group was primed at week 0 with 100 ug SARS-CoV-2 spike protein immunogen with 375 ug of SMNP adjuvant. The dose escalation group was primed using seven increasing doses, with 0.2, 0.43, 1.16, 3.15, 8.56, 23.3, and 63.2 μgs of SARS-CoV-2 spike protein in SMNP adjuvant being administered on days 0, 2, 4, 6, 8, 10, and 12 respectively. Both groups were then boosted with 100 μg SARS-CoV-2 spike protein and 375 ug SMNP at week 10. For the present study, the same eight rhesus macaques were all immunized in the same manner with 100 μg MERS-CoV spike protein in 375 ug SMNP at week 38, and 100 μg of an equally divided cocktail of the spike proteins of MERS-CoV, HCoV-HKU1 and HCoV-OC43 in 375 ug SMNP at week 46. Multiple times during the study, EDTA blood was collected for PBMC and plasma isolation using CPT tubes, and serum was isolated in serum collection tubes and frozen for later use.

### Plasmid Construction

Expression plasmids of the spike (S) protein ectodomain proteins were constructed by cloning the S-protein encoding genes synthesized by GeneArt (Life Technologies) from SARS-CoV-1 (residues 1-1190; GenBank: AAP13567), SARS-CoV-2 (residues 1-1208; GenBank: MN908947), HCoV-HKU1 (residues 1-1295; GenBank: YP_173238.1), HCoV-OC43 (residues 1-1300; GenBank: AAX84792.1), MERS-CoV (residues 1-1291; GenBank: APB87319.1), HCoV-229E (residues 1-1110; GenBank: NP_073551.1) and HCoV-NL63 (residues 1-1291; GenBank: YP_003767.1) into the phCMV3 vector (Genlantis Cat# P003300) using Gibson assembly [New England Biolabs (NEB), E2621L]. To produce soluble stabilized trimeric spike proteins, the furin cleavage site of each S protein was replaced with a “GSAS“ linker (in SARS-CoV-2 residues 682–685, in SARS-CoV-1 residues 664–667, in HCoV-HKU1 residues 756-760, in HCoV-OC43 residues 762–766, in MERS-CoV residues 748–751, in HCoV-229E residues 564–567 and in HCoV-NL63 residues 745–748). The following residues were changed to prolines to maintain the trimer’s pre-fusion conformation; SARS-CoV-1: K968+V969, SARS-CoV-2: K986+V987, MERS-CoV: V1060+L1061, HCoV-HKU1: A1071+L1072, HCoV-OC43: A1078+L1079, HCoV-NL63: S1052+I1053, HCoV-229E: T871+I872 (Hsieh et al., 2020; Pallesen et al., 2017). Additionally, a trimerization T4 fibrin motif was added to the C-terminus of each S protein, followed by a 6x HisTag and AviTag (Avidity) separated by GS-linkers to facilitate purification and biotinylation of the proteins. To generate the SARS-CoV-2 soluble monomeric RBD expression plasmid, S-protein residues 320 to 527 were cloned into the phCMV3 vector (Genlantis Cat# P003300) with the Tissue Plasminogen Activator leader sequence added to the N-terminus and 6x HisTag and AviTag separated by GS-linkers added to the C-terminus.

### Soluble Spike and RBD protein expression and purification

Freestyle293F cells (Thermo Fisher R79007) were used to transfect plasmids encoding the human coronavirus soluble S ectodomain and RBD proteins. 350μg of each respective coronavirus protein 100μg were diluted into 40 mL of Transfectagro™ (Corning 40-300-CV). 100ug of BirA coenzyme encoding plasmid (Avidity) was added for proteins used as biotinylated probes in B cell sorting, BLI analysis, or ELISA assays. This mixture was then filtered using 0.22 μm Steriflip™ Sterile Disposable Vacuum Filter Units (MilliporeSigma™ SCGP00525) before gently mixing in 1.6 mL of 40 K polyethylenimine (Polysciences 24765-1) (1 mg/mL) and letting the resulting solution incubate for 30 minutes at room temperature. Following incubation, the mixture was added to 1 L of Freestyle293F cells at 1 million cells mL^-1^. After four days, cultures were spun down for 15 minutes at 2500 x g, the supernatant was filtered into glass bottles with 0.2 μm membrane filters (Fisher Scientific 564-0020), and then the cultures were stored at 4°C for purification. Protein constructs containing a His-tag were purified by slowly passing the culture supernatant through HisPur Ni-NTA Resin (Thermo Fisher 88221) beads in columns overnight. The next day, beads were washed with three bead volumes of wash buffer (25mM Imidazole, pH 7.4), and then finally eluted with 25ml (250mM Imidazole, pH 7.4). Protein constructs without a His-tag were purified with GNL beads (Vector Labs AL-1243-5), washed with 3 column volumes of PBS, and eluted with 1M Methyl α-D-mannopyranoside (Sigma M6882). Using Amicon® 100 kDa for spike proteins or 10 kDa Ultra-15 Centrifugal Filter Units for RBD proteins (Merck Millipore UFC9100 & UFC9010), the proteins were buffer exchanged in PBS and then concentrated down to 500µl for size exclusion chromatography using a Superdex 200 Increase 10/300 GL column (Sigma-Aldrich GE28-9909-44). Specific fractions from the chromatography run were mixed and concentrated before being used.

### mAb expression and purification

Expi293F cells (Thermo Fisher Scientific A14527) were used for the transfection of mAbs. HC and LC variable regions of select mAbs were cloned into plasmid expression vectors with their respective constant regions. Antibody sequences have been deposited in GenBank under accession nos. PV206838-PV206933. Per transfection, 16μg of both heavy and light chain encoding plasmid were diluted in 4ml Opti-MEM (Gibco 31985070) and then gently mixed with 32 μl FectoPRO transfection reagent (116-040; Polyplus). After a 10-minute incubation at room temperature, the solution was added to 40 mL of cells at 2.8 million cells ml^-1^. 24 hours after transfection, cells were fed with 400 μl sodium valproic acid (300mM) and 360 μl 45% D-(+)-Glucose Solution (Sigma G8769-100ML) to boost transfection yield. After five days, the cell cultures were spun down for 15 minutes at 2500 x g. The supernatants were collected and filtered using 0.22μm Steriflip™ Sterile Disposable Vacuum Filter Units (MilliporeSigma™ SCGP00525), and then stored at 4°C for purification. For purification, mAbs were rotated overnight with 0.25ml of Praesto Protein A Affinity Chromatography Resin (Purolite PR00300-164) and 0.25ml Protein G Sepharose (Cytiva GE17-0618-01). The next day, beads were transferred to an Econo-Pac column (Bio-Rad Laboratories 7321010), washed with 2 column volumes of PBS, and then eluted with 10 ml 0.2M citric acid (pH 3). After elution, the solution’s pH was adjusted using 1ml 2M Tris base (pH 9). Using Amicon® 30 kDa Ultra-15 Centrifugal Filter Units (Merck Millipore UFC9030), the mAbs were buffer exchanged into PBS and then concentrated before use. mAbs used for protection experiments also underwent further size exclusion chromatography using a Superdex 200 Increase 10/300 GL column (Sigma-Aldrich GE28-9909-44). Specific fractions from the chromatography run were mixed and concentrated before being used.

### Production of coronavirus pseudoviruses

2.5µg of plasmid encoding for the spike protein of each virus, with the endoplasmic reticulum retrieval signal removed, was co-transfected with 10µg MLV (murine leukemia virus)-CMV (cytomegalovirus) luciferase and 12.5µg MLV Gag/Pol plasmids into HEK-293T cells (ATCC CRL-3216) using Lipofectamine 2000 (Thermo Fisher 11668019) following the manufacturer’s protocol. After 16 hours, cell media was removed and replaced with warmed media (DMEM (Corning 10-017-CV) with 10% FBS (Omega Scientific NC0471611), 1% L-glutamine (Corning 25-005-CI), and 1% penicillin-streptomycin (Corning 30-002-CI)). After 48 hours, the cell supernatant was collected, filtered with 0.22 μm Steriflip™ Sterile Disposable Vacuum Filter Units (MilliporeSigma™ SCGP00525), and stored at −80°C.

### Pseudovirus neutralization assay

Target cells were created by using a lentivirus system to transduce hACE2 into HeLa cells (ATCC CCL-2). Cells with the highest and most stable hACE2 expression were isolated and cultured for use. For the neutralization assay, mAbs were diluted in cell media (DMEM (Corning 10-017-CV) with 10% FBS (Omega Scientific NC0471611), 1% L-glutamine (Corning 25-005-CI) to a starting concentration of 600 ug/ml and serum samples were diluted in cell media to a starting dilution of 1:15. Both mAbs and serum were then serially diluted. 25 µl of diluted mAbs and serum were added to assay plates (Corning 3688). 25 µl of pseudovirus was added to diluted antibodies and then the plates were incubated at 37°C. After 1 hour, 50 µl of HeLa hACE2 cells at 2x10^5^ cells ml^-1^ with 20 μg ml^-1^ DEAE-dextran (Sigma-Aldrich 93556-1G) was added to each well. Plates were then incubated at 37°C for 48 hours. After incubation, supernatant was removed from plates and 60ul of a 1:10 ratio of luciferase lysis buffer (25 mM Gly-Gly, pH 7.8, 15 mM MgSO_4_, 4 mM EGTA, 1% Triton X-100) and Bright-Glo (Promega Corporation E2620) was used to lyse the hACE2 cells and initiate fluorescence. Amount of luminescence was quantified using a luminometer. All samples were tested in duplicate. The following equation was used to determine percent neutralization: Percentage neutralization = 100 × (1 − ((RLU of sample)−(Average RLU of CC))/((Average RLU of VC)−(Average RLU of CC))). Calculations and graphing were completed in Prism 8 (Graph Pad Software). IC50 was determined by fitting the luminometer readings to a non-linear regression and then measuring concentration of mAb at 50% neutralization. ID_50_ was determined by fitting the luminometer readings to a non-linear regression and then measuring the dilution factor of serum at 50% neutralization.

### Fab isolation from poly-IgG

For polyclonal serum, poly-IgG was first isolated by rotating 1ml of serum with 0.25 ml of Praesto Protein A Affinity Chromatography Resin (Purolite PR00300-164) and 0.25 ml Protein G Sepharose (Cytiva GE17-0618-01), and 9 ml PBS. The next day, beads were transferred to an Econo-Pac column (Bio-Rad Laboratories 7321010), washed with 2 column volumes of PBS, and then eluted with 10 ml 0.2M citric acid (pH 3). After elution, the solution’s pH was adjusted using 1 ml 2M Tris base (pH 9). Using Amicon® 30 kDa Ultra-15 Centrifugal Filter Units (Merck Millipore UFC9030), the resulting poly-IgG Abs were buffer exchanged into PBS and then concentrated. Polyclonal antibodies and mAbs were then digested using Pierce™ Fab Preparation Kit (Thermo Fisher 44985) following manufacturer instructions. After digestion, Amicon® 10 kDa Ultra-15 Centrifugal Filter Units (Merck Millipore UFC9010) were used to buffer exchange proteins into PBS and concentrate to 0.5 mL. Concentrated Fabs underwent size exclusion chromatography using Superdex 200 Increase 10/300 GL column (Sigma-Aldrich GE28-9909-44). Specific fractions from the chromatography run were mixed and concentrated before being used.

### Enzyme-linked immunosorbent assay (ELISA)

96-well half well plates were coated with 2 µg/mL streptavidin (Jackson ImmunoResearch Laboratories, # 016-000-084) in PBS overnight at 4°C. Plates were then washed three times with PBS + 0.05% Tween20 (PBST) and blocked with 3% bovine serum albumin (BSA) in PBS for 2 h at room temperature (RT). After removing the blocking solution, biotinylated proteins were added in PBST + 1% BSA and incubated for 1.5 h at RT. Spike or RBD proteins were added at 2 μg/ml; S2 peptides were added at 16 µg/ml. After washing the plates three times with PBST, samples were added and incubated for 1.5 h at RT. After three washes with PBST, the secondary antibody (AffiniPure Goat anti-human IgG Fc fragment specific, Jackson ImmunoResearch Laboratories, #109-055-008) was added and incubated for 1h at RT. Three washes with PBST later, phosphatase substrate (Sigma-Aldrich, # S0942) was added into each well and incubated at RT for 10 min. Absorbance was measured at 405 nm using a using VersaMax microplate reader (Molecular Devices) and analyzed using SoftMax version 5.4 (Molecular Devices).

Biotinylated spike or RBD proteins were produced in house as described above; N-terminal biotinylated S2 stem helix peptides (SARS-CoV-2 residues 1140-1164) were synthesized at GenScript as follows:

SARS-CoV-1/2 (PLQPELDSFKEELDKYFKNHTSPDV), MERS-CoV (PLLGNSTGIDFQDELDEFFKNVSTSIP), HCoV-HKU1 (HSVPKLSDFESELSHWFKNQTSIAP), HCoV-OC43 (TSIPNLPDFKEELDQWFKNQTSVAP), HCoV-229E (TIVPEYIDVNKTLQELSYKLPNYTV), and HCoV-NL63 (TVIPDYVDVNKTLQEFAQNLPKYVK).

### BioLayer Interferometry (BLI) competition assay

Samples were analyzed with an Octet HTX instrument using Anti-SA biosensors (Sartorius 18-5019). Soluble spike proteins conjugated with biotin were diluted to 100 nM, and mAbs were diluted to 10 µg/ml in Octet buffer (1X PBS with 0.1% Tween20). For competition experiments, hydrated biosensors captured spike protein for 300 s, then baseline was determined by transferring probe to Octet buffer for 30 s. Next, probes were placed in saturating antibody for 200 s, transferred to buffer for 30 s, and then placed in buffer containing competing antibody for 200 s. Competition was determined by subtracting the response measurement after the competing antibody was bound by the response when the saturating antibody was bound. The larger the difference in response between the competing and saturating antibody, the less competition between the two antibodies.

### BioLayer Interferometry (BLI) S2 alanine scanning assay

Wild type S2 stem helix peptides, as well as single alanine mutation-bearing 25-mer stem helix peptides, were produced by GenScript. Samples were analyzed with an Octet HTX instrument using Anti-SA biosensors (Sartorius 18-5019). S2 peptides conjugated with biotin were diluted to 1 μg/ml, and mAbs were diluted to 10 µg/ml in Octet buffer (PBS with 0.1% Tween20). Hydrated biosensors were first used to capture S2 stem helix peptides for 60 seconds, and then the probes were transferred to Octet buffer to determine a baseline measurement. Probes were then transferred into monoclonal antibody solutions for 120 s to determine maximum binding responses, and finally tested for dissociation in Octet buffer for 240 s.

### Negative stain electron microscopy

Spike constructs (HexaPro-GSAS D614G, HexaPro-GSAS Mut2 D614G, HexaPro-GSAS Mut7 D614G, and HexaPro Mut7 D614G) for EM studies were generated and expressed as described previously (Bangaru et al., 2022). Constructs with intact S1/S2 cleavage site were co-expressed with furin. All constructs contain six stabilizing proline (HexaPro) substitutions at positions 817, 892, 899, 942, 986, and 987. Additional cysteine substitutions at 383 and 985 were included for Mut2, and 705 and 883 for Mut7. All constructs for polyclonal Fab complexes contain the D614G mutation and those labeled with GSAS have the S1/S2 furin cleavage site modified to 682-GSAS-685. For the polyclonal complexes, a mix of HexaPro-GSAS D614G, HexaPro-GSAS Mut2 D614G, HexaPro-GSAS Mut7 D614G, and HexaPro Mut7 D614G was incubated with a 25-fold molar excess of polyclonal fabs at room temperature for one hour. A Superose 6 Increase 10/300 GL column in an AKTA Pure system was used to isolate complexes from unbound Fab. For the monoclonal complexes, Fabs were complexed with HexaPro Mut7 at a 9-fold molar excess relative to spike for one hour at room temperature without SEC purification. Complexes were concentrated to ∼0.03 mg/ml the same day using 1X TBS pH 7.4 and applied to glow-discharged copper mesh grids before staining with 2% uranyl formate for 90 seconds. A 4K x 4K Thermo Scientific Ceta 16M camera on a Talos F200C (200keV, 73kx mag), TIETZ 4K x 4K camera on a FEI Tecnai Spirit (120keV, 52kx mag), and Thermo Fisher Falcon 4i Direct Electron Detector 4K x 4K camera on a Thermo Fisher Glacios (200 keV, 73k x mag) were used to collect data. Automated data collection on the Talos and Spirit was done with Leginon (Suloway et al., 2005), and micrographs were stored in the Appion database(Lander et al., 2009). Particles were picked using DoGpicker (Voss et al., 2009) and further processed in Relion 3.0 (Zivanov et al., 2018). Week 40 datasets were also independently processed with the inclusion of all spike particles (intact and dissociated trimers) and 3D classification was carried out in Relion 3.0 using a mask-based classification approach. A spherical mask was placed around the spike ectodomain with the exclusion of S2 SH region to prevent any bias in alignment by the dominant S2 SH pAbs. Micrographs from Glacios were completely processed in Relion 3.1 (Scheres, 2012). 3D models were segmented using UCSF Chimera (Pettersen et al., 2004).

### B cell sorting

Frozen cells harvested from NHP inguinal lymph nodes were flash-thawed in 10mL recovery medium (RPMI 1640 medium containing 50% FBS). Cells were centrifuged for 5 min at 500 rcf and resuspended in ice-cold FACS buffer (PBS+2% FBS). The sample was counted, spun down, and resuspended to a target concentration of 1x10^7^ cells/ml.

100 µl of cell solution were transferred to a new plate or falcon tube. 5 µl of anti-Rh Fc receptor binding inhibitor (eBioScience, #14-9165-42) was added per sample and incubated for 15 min on ice. After incubation, the cells were spun down and incubated with staining mix in Brilliant Stain Buffer (Invitrogen, #00-4409-42) for 30 min on ice in the dark. Antibodies used were: CD3 (APC Cy7, BD Pharmingen, clone SP34-2, #557757), CD4 (APC-Cy7, Biolegend, clone OKT-4, #317418), CD8 (APC-Cy7, Biolegend, clone RPA-T8, #557760), CD14 (APC-H7, BD Pharmingen, clone M5E2, #561384), CD19 (PerCP-Cy5.5, Biolegend, clone HIB19, #302230), CD20 (PerCP-Cy5.5, Biolegend, clone 2H7, #302326), IgG (BV786, BD Horizon, clone G18-145, #564230) and IgM (PE, Biolegend, clone MHM-88, #314508). After 30 minutes, protein probes were added. Probes were prepared as follows: Streptavidin-AF488 (Thermo Fisher S32354) and streptavidin-BV421 (BD Biosciences 563259) were coupled to BirA biotinylated SARS-CoV-2 spike protein separately at a molar ratio of 3:4 streptavidin:antigen-biotin. After 30 min incubation at RT, the conjugated spike proteins were stored on ice. After adding 1.6 µg of SARS-CoV-2 spike (AF488) and 0.55 ug SARS-CoV-2 spike (BV421) to the cells, the mix was incubated for 15 min on ice in the dark. Afterwards the cells were spun down, resuspended in FVS510 Live/Dead stain (Thermo Fisher Scientific, #L34966) as 1:1000 dilution in FACS buffer, and incubated for 15 min on ice in the dark. Subsequently, the cells were washed once with ice cold FACS buffer, resuspended in 300µL ice cold FACS buffer, and filtered through 70 um nylon mesh FACS tube caps (Fisher Scientific, #08-771-23). Before cell sorting on a BD FACSMelody sorter (BRV 9 Color Plate 4way), a well of a 96-well-plate was prepared by coating the side of the wells with 200 µL 100% FBS. Excess FBS was removed and 20 µL of 100% FBS was added at the bottom of the well. CD19+ CD20+ CD3- CD4- CD8- CD14- IgM- IgG+ live singlet cells that were positive for both SARS-CoV-2 spike (AF488) and SARS-CoV-2 spike (BV421) were sorted into the prepared well to a maximum of 20000 cells. After sorting, PBS was added to the used wells to dilute the FBS concentration to below 10%. The plates were covered with aluminum plate covers and spun down at 2500 RPM for 2 min at 4°C. Following this step, the workflow followed the 10x Genomics’ ‘Next GEM Single Cell 5’ Kit’ manufacturer’s instructions for sample preparation and library construction. Custom primers were used for step ‘V(D)J Amplification from cDNA’:

#### B Cell Mix 1 Rhesus Macaque

IGHA-R1 CAGGGCACAGCCACATCCT

IGHD-R1 GTGCTTGACGGTGCATTTGTA

IGHE-R1 AAGGTTTTGTTGACCTCTTTGTCT

IGHG-R1 TTGTCCACCTTGGTGTTGCT

IGHM-R1 CATGACGTCCTTGGAAGCCA

IGKC-R1 TCTGGTAGTCTGTGCTGCTCAG

IGLC-R1 TGTGGGACTTCCACTG

R1-Forward

AATGATACGGCGACCACCGAGATCTACACTCTTTCCCTACACGACGCTC

#### B Cell Mix 2 Rhesus Macaque

IGHA-R2 CGGGAAGTTTATGACGGTCA

IGHD-R2 GGGTTGTACCCAGTTATCAAGCAT

IGHE-R2 GCGTCCCAGGTCACCATCAC

IGHG-R2 CCCTGAGGACTGTAGGACAGC

IGHM-R2 GCCACTTCGTTTGTATCCAA

IGKC-R2 GACACCATCCACCTTCCACTTT

IGLC-R2 TAGCTGCTGGCCGC

R2-Forward AATGATACGGCGACCACCGAGATCT

### Sequence characteristics of isolated mAbs

Analysis of the heavy chain (HC) variable region (VH) and light chain (LC) variable region (VK/VL) gene usage, number of V region somatic hypermutations (SHM), complementary determining region 3 (CDR3) sequences, and CDR3 lengths for each antibody was done using the rhesus macaque (*Macaca mulatta*) germline database from IMGT (https://www.imgt.org/IMGT_vquest/analysis).

### B cell profiling

Frozen PBMC samples were processed as described for B cell sorting. The cells were stained with the following antibodies: CD8 (clone RPA-T8, BD Pharmingen #557760), CD14 (M5E2, BD Pharmingen #561384), IgM (MHM-88, Biolegend #314508), CD4 (OKT-4, Biolegend #317411), CXCR5 (Mu5UBEE, ThermoFisher #25-9185-41), CD38 (AT-1, Stem Cell Technologies #60131FI), CD20 (2H7, Biolegend #302326), CD27 (O323, Biolegend #302825), PD1 (EH12.2H7, Biolegend #329949), IgG (G18-145, BD Horizon #564230), CD71 (L01.1, BD Horizon #749821). Protein probes used were prepared by conjugating BirA biotinylated proteins (MERS-CoV spike, HCoV-OC43 spike, SARS-CoV-2 spike, and SARS-CoV-2 RBD) to streptavidin-bound fluorophores as described for B cell sorting. After staining the cells with antibody mix, protein probes and Live-Dead stain, the cells were washed once with ice cold FACS buffer, resuspended in 300 µL ice cold FACS buffer, and filtered through 70 um nylon mesh FACS tube caps (Fisher Scientific, #08-771-23). Subsequently, the samples were processed on a Bigfoot Spectral Cell Sorter (Invitrogen). Cytometry data were analyzed using FlowJo version 10 (TreeStar).

### Monoclonal Fab purification

Following manufacturer instructions, the Pierce™ Fab Preparation Kit (Thermo Fisher 44985) was used to isolate Fab proteins from previously expressed CHM-16 and CHM-27 IgG protein. Fabs were then buffer exchanged into Tris-buffered saline (TBS) and concentrated using Amicon® 10 kDa Ultra-15 Centrifugal Filter Units (Merck Millipore UFC9010). Concentrated proteins were run through further size exclusion chromatography using a Superdex 200 Increase 10/300 GL column (Sigma-Aldrich GE28-9909-44). Specific fractions from the chromatography run were mixed and concentrated before use.

### Crystallization and structural determination

CHM-16 Fab (11 mg/ml) with 10× (molar excess) SARS-CoV-2 stem peptide (1140-PLQPELDSFKEELDKYFKNHTSPDV-1164) as well as CHM-27 Fab (13 mg/ml) with 9× (molar excess) MERS-CoV stem peptide (1221-PLLGNSTGIDFQDELDEFFKNVSTSIP-1247) were screened for crystallization using the 384 conditions of the JCSG Core Suite (Qiagen) on our robotic CrystalMation system (Rigaku) at Scripps Research. Crystallization trials were set up by the vapor diffusion method in sitting drops containing 0.1 μl of protein and 0.1 μl of reservoir solution. Crystals of CHM-16 with SARS-CoV-2 stem peptide were grown in drops containing 0.1 M sodium cacodylate pH 6.5, and 1 M sodium citrate at 20°C. Crystals appeared on day 7 and were harvested on day 18 by soaking in reservoir solution supplemented with 15% (v/v) glycerol as cryoprotectant. Optimized crystals of CHM-27 in complex with MERS-CoV S2 stem peptide were grown in drops containing 0.17 M sodium acetate, 0.085 M Tris at pH 8.5, 15% (v/v) glycerol, and 25.5% (w/v) polyethylene glycol 4000 at 20°C. Crystals of CHM-27-peptide complex appeared on day 3, were harvested on day 14 by soaking in reservoir solution supplemented with 20% (v/v) ethylene glycol as cryoprotectant. The crystals were then flash-cooled and stored in liquid nitrogen until data collection. Diffraction data of CHM-16-peptide complex and CHM-27-peptide complex were collected at cryogenic temperature (100 K) at National Synchrotron Light Source II (NSLS-II) on beamlines 17-ID-1 and 17-ID-2, with beam wavelengths of 0.92010 Å and 0.97933 Å, respectively. Diffraction data were processed with HKL2000 (PubMed: 27754618) and AutoPROC (Vonrhein et al., 2011). Structures were solved by molecular replacement using PHASER (McCoy et al., 2007) and CC25.106 with SARS-CoV-2-SH peptide as initial model (PDB: 8DGU). Iterative model building and refinement were carried out in COOT(Emsley et al., 2010) and PHENIX(Adams et al., 2010), respectively. Epitope and paratope residues, as well as their interactions, were identified by accessing PISA at the European Bioinformatics Institute (http://www.ebi.ac.uk/pdbe/prot_int/pistart.html)(Krissinel and Henrick, 2007) .

### Murine viral challenge

Groups of K18 mice expressing human ACE2 receptors were treated intraperitoneally (i.p.) with 300 µg per animal of S2 SH mAb, positive, or negative control mAbs. 12 hours after mAb infusion, serum was collected from all animals. Animals that did not show titers of S2 SH, positive or negative control antibodies, respectively, were excluded from the study (CHM-16: n=2, CHM-27: n=3, CC12.1: n=0. SMZAb1: n=0). Mice showing titers of the treatment antibodies were challenged with 2 × 10^4^ focus forming units (FFU) of SARS-CoV-2 (USA-WA1/2020) virus. Animals were weighed daily and euthanized at day 6 post challenge. Lung tissue of all animals was collected for further analysis.

### Lung virus titer analysis

Lungs were resuspended in 0.5 mL of 2% FBS complete DMEM and homogenized. Lung lysate was added to HelaACE2 cells and incubated overnight at 37°C. Cells were then fixed with 4% PFA for 1hr. Plates were then developed with a primary stain of diluted convalescent plasma from a COVID-19-positive patient diluted in 1X Perm/Wash solution. Secondary stain was an anti-human FC conjugated with HRP (AffiniPure Goat anti-human IgG Fc fragment specific, Jackson ImmunoResearch Laboratories, #109-055-008). The final development step of the plates required adding ‘TruBlue’ Peroxidase Substrate (VWR, #95059-468) and then counting for focus forming units (FFU).

### Lung viral RNA analysis

Viral RNA was isolated from lung tissue and subsequently amplified and quantified in a reverse transcription (RT) qPCR. Lung tissue was extracted at day 6 after infection and placed in 0.5 ml of TRIzol reagent (Invitrogen). The samples were then homogenized using Bead Ruptor 12 (Omni International). Tissue homogenates were then spun down, and the supernatant was added to an RNA purification column (Qiagen). Purified RNA was eluted in 60 μl of deoxyribonuclease-, ribonuclease-, endotoxin-free molecular biology–grade water (Millipore). RNA was then subjected to RT and qPCR using the Centers for Disease Control and Prevention’s N1 (nucleocapsid) primer sets (forward, 5′-GACCCCAAAATCAGCGAAAT-3′; reverse, 5′-TCTGGTTACTGCCAGTTGAATCTG-3′) and a fluorescently labeled (FAM) probe (5′-FAM-ACCCCGCATTACGTTTGGTGGACC-BHQ1-3′) (Integrated DNA Technologies) on a Bio-Rad CFX96 real-time instrument. For quantification, a standard curve was generated by diluting 8.5 × 105 PFU RNA equivalents of SARS-CoV-2. Every run used 11 5-fold serial dilutions of the standard. SARS-CoV-2–negative mouse lung RNA and no templates were both included as negative controls for the extraction step as well as the qPCR.

## QUANTIFICATION AND STATISTICAL ANALYSIS

Statistical analyses were performed using GraphPad Prism 8 (GraphPad), San Diego, California, USA. Group data comparison for B cell profiling and animal challenge experiments was performed using one-way ANOVA or two-way ANOVA with Holm-Šídák multiple comparison test. Data were considered statistically significant at p < 0.05. Statistical details of experiments can be found in the Method Details section and figure legends.

## Supplemental information

**Figure S1.**
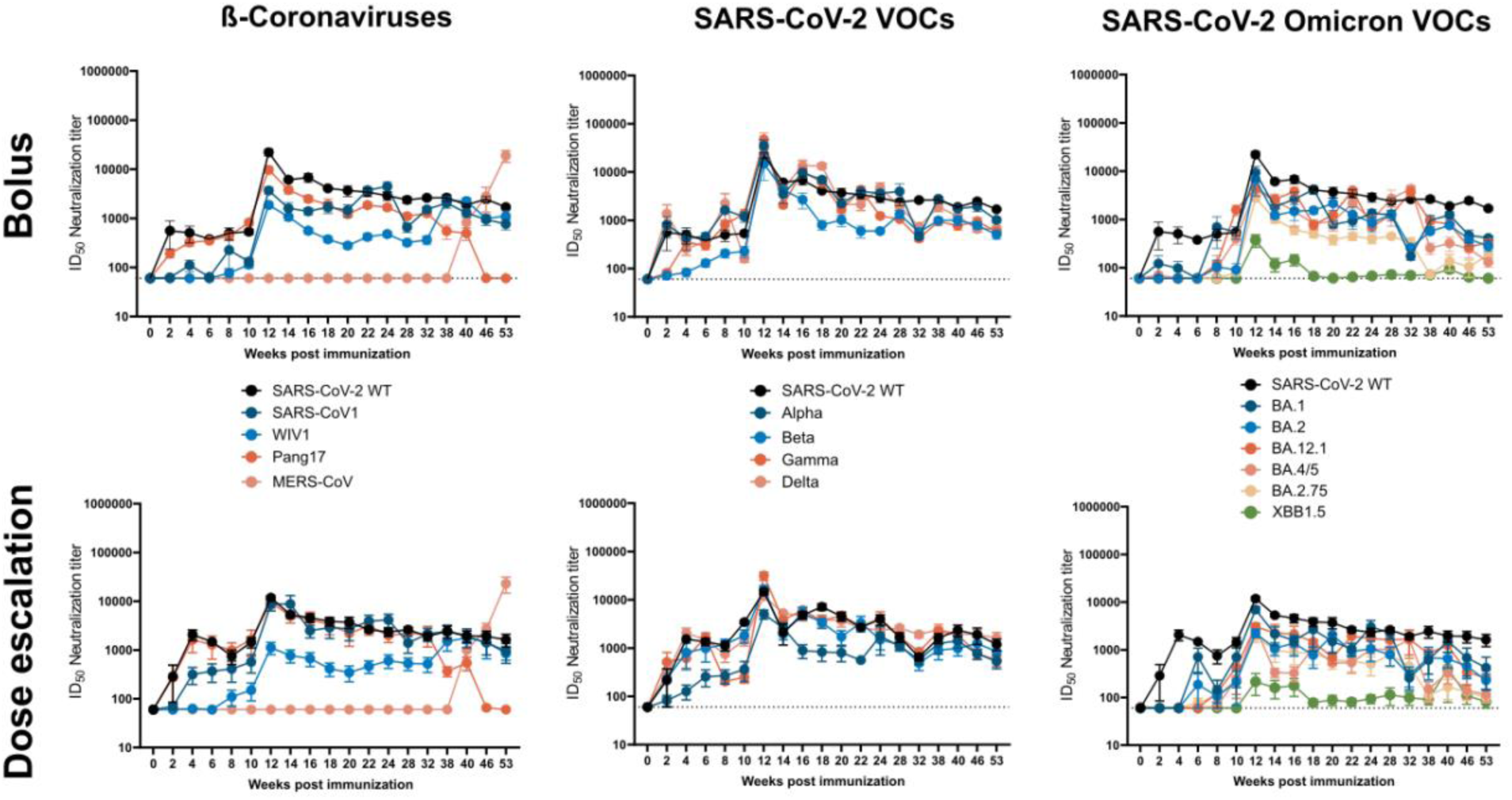
Serum pseudovirus neutralization titers as a function of time (cf. Fig 1) for bolus and dose escalation immunization regimes. Shown are serum ID_50_ neutralization titers for the indicated pseudoviruses for the bolus group and the dose escalation group (n=4 each). Data points are shown as mean ± SEM.

**Figure S2.**
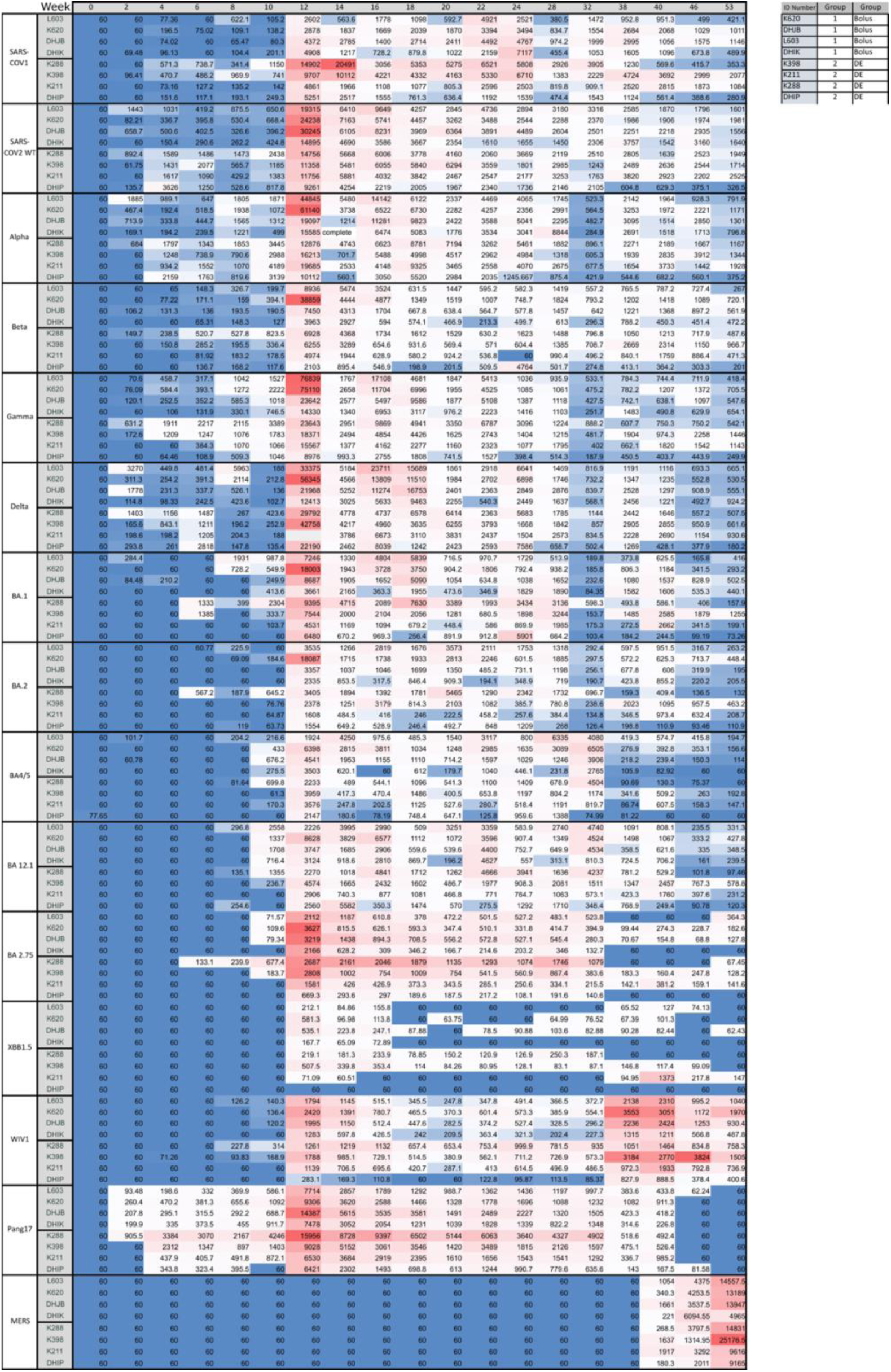
Serum pseudovirus neutralization titers as a function of time by individual animal. Shown are serum ID_50_ neutralization titers for the indicated pseudoviruses for each individual macaque. Data points are shown as mean ± SEM. cf.

**Figure S3.**
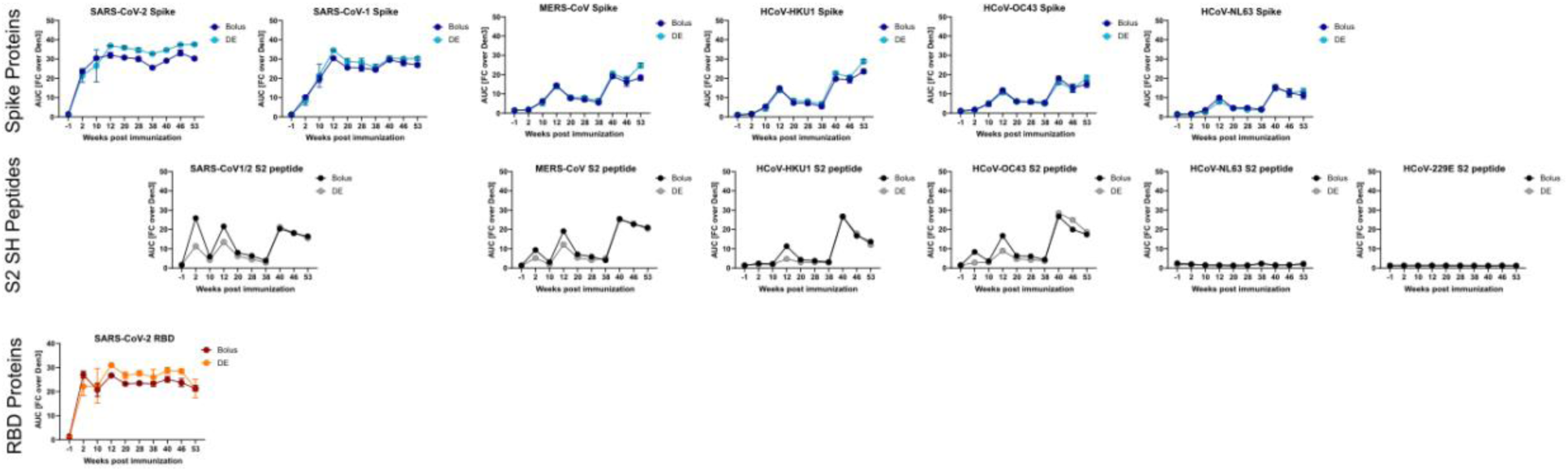
Serum antibody binding to spike proteins and peptides as a function of time for bolus and dose escalation regimes. NHP serum binding to spike proteins (upper panel), S2 stem helix peptides (middle panel). and receptor binding domains (lower panel) of the indicated viruses was measured by ELISA at the indicated time points. Data are shown as area under curve (AUC) as fold change over negative control, Den3. Data are shown as mean ± SEM. N=4 for both groups. cf.

**Figure S4.**
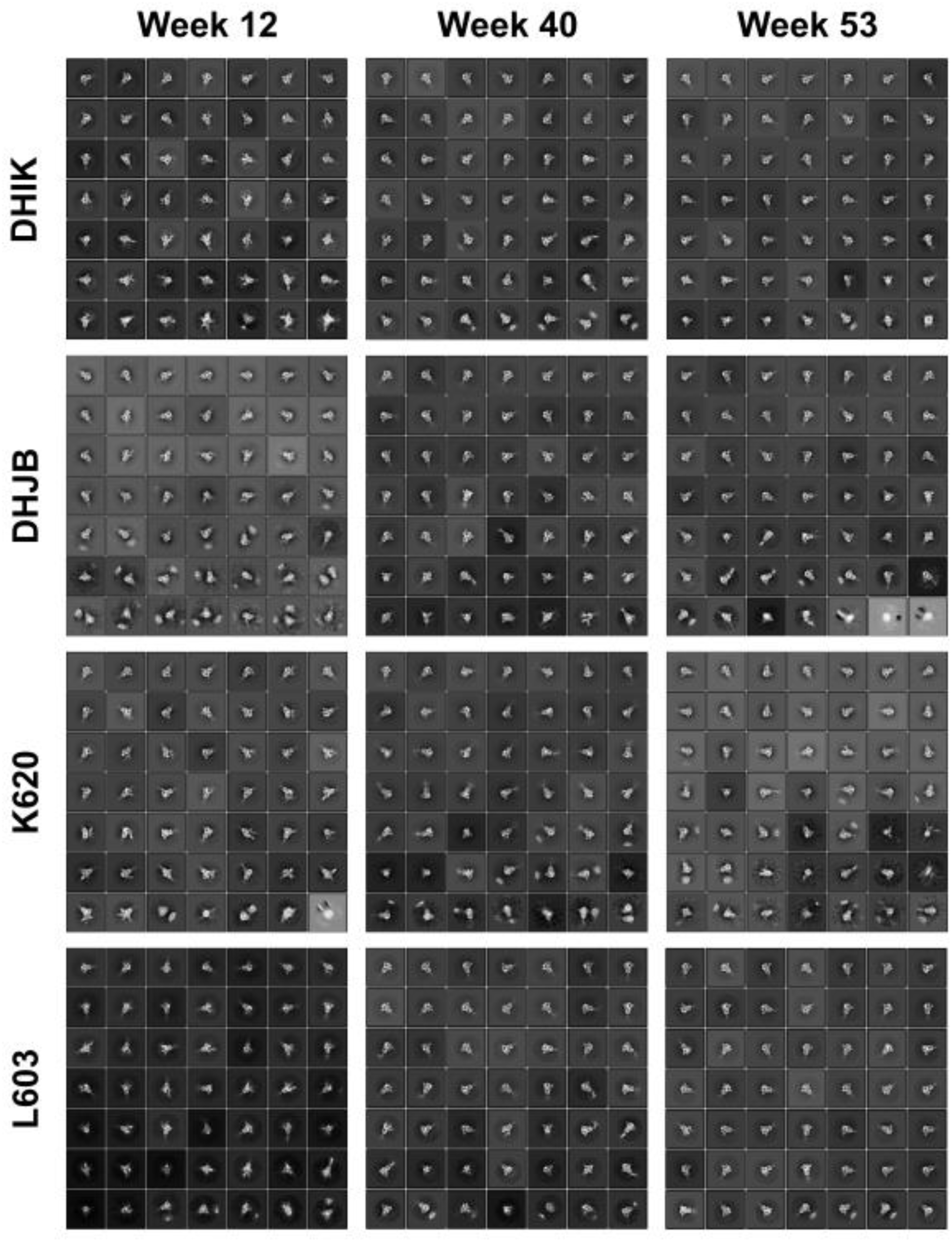
Epitope mapping of S2 SH bnAbs from negative stain Electron Microscopy (ns-EM). 2D class averages of Fabs isolated from the plasma of 4 animals (K620, L603, DHIK and DHJB) are shown in complex with SARS-CoV-2 (Wuhan) spike protein.

**Figure S5.**
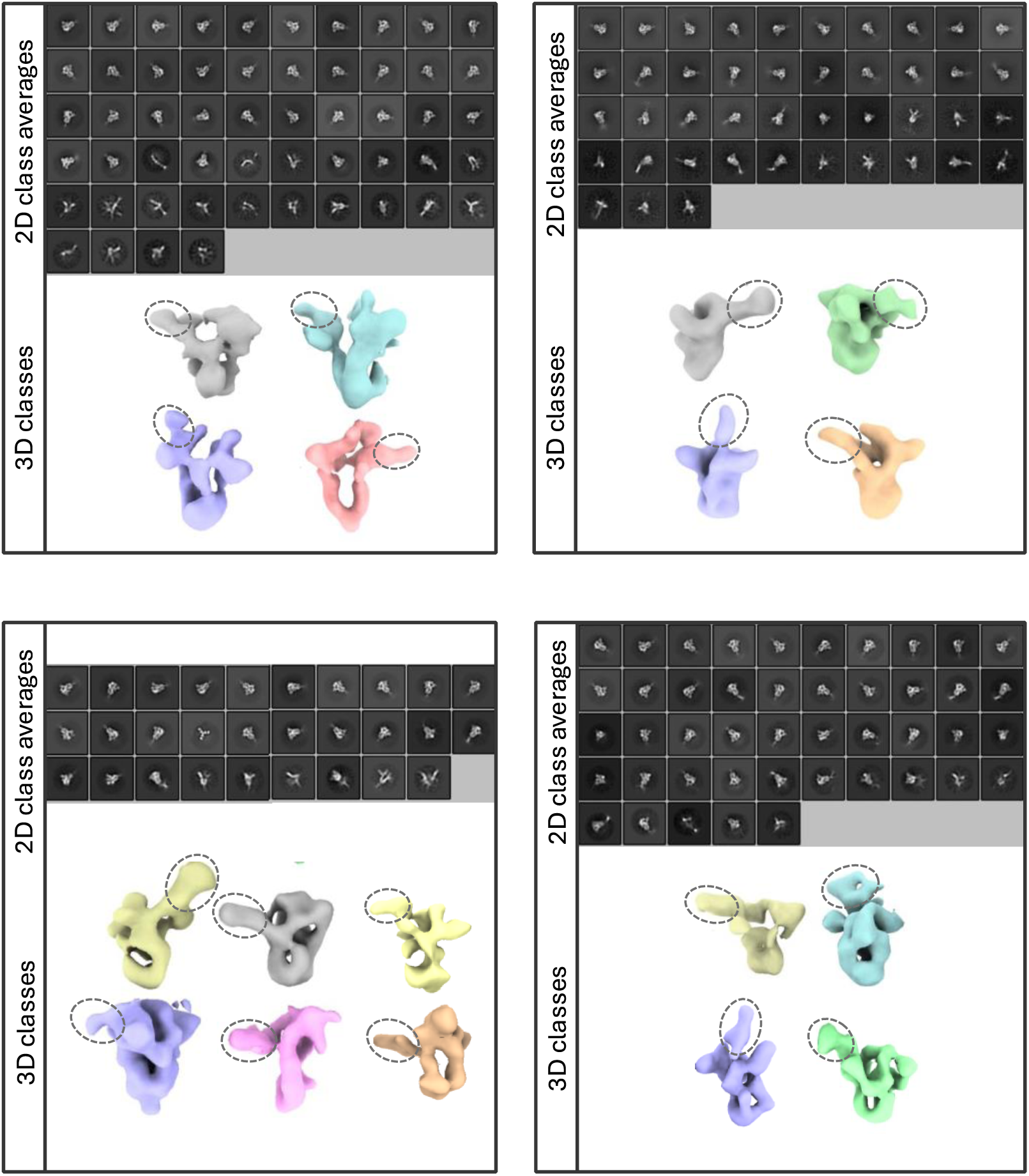
Ns-EMPEM analysis of responses from week 40 sera with the inclusion of dissociated spikes in data processing. 2D class averages and 3D classes generated with mask-based 3D classification (excludes S2 stem-helix region) show the presence of dissociated spikes with Fab densities (dotted circles)

**Figure S6.**
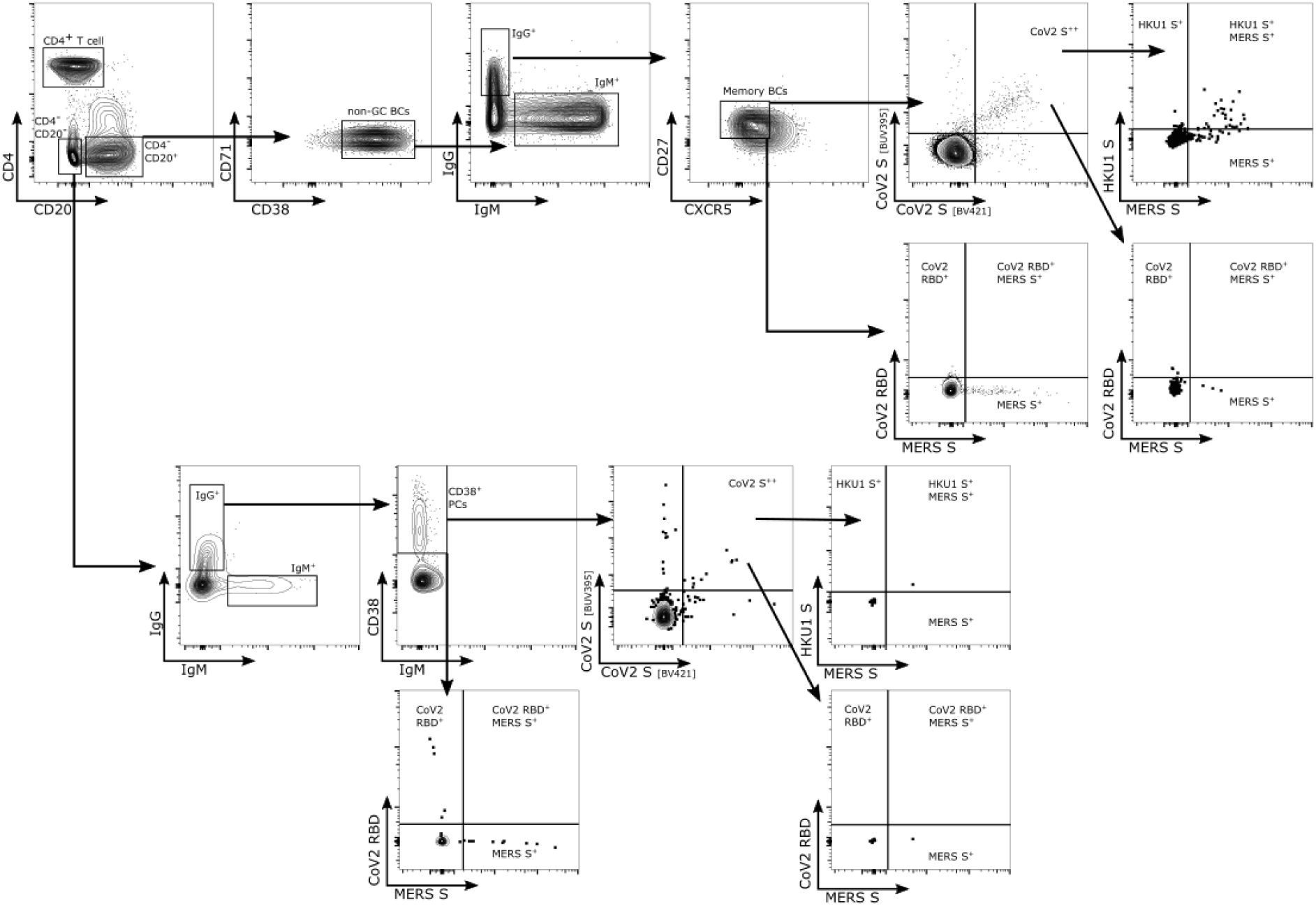
Gating strategy for PBMC analysis. PBMCs of select animals (3 per group) were analyzed using the presented gating strategy. Data are representative of all samples analyzed.

**Figure S7.**
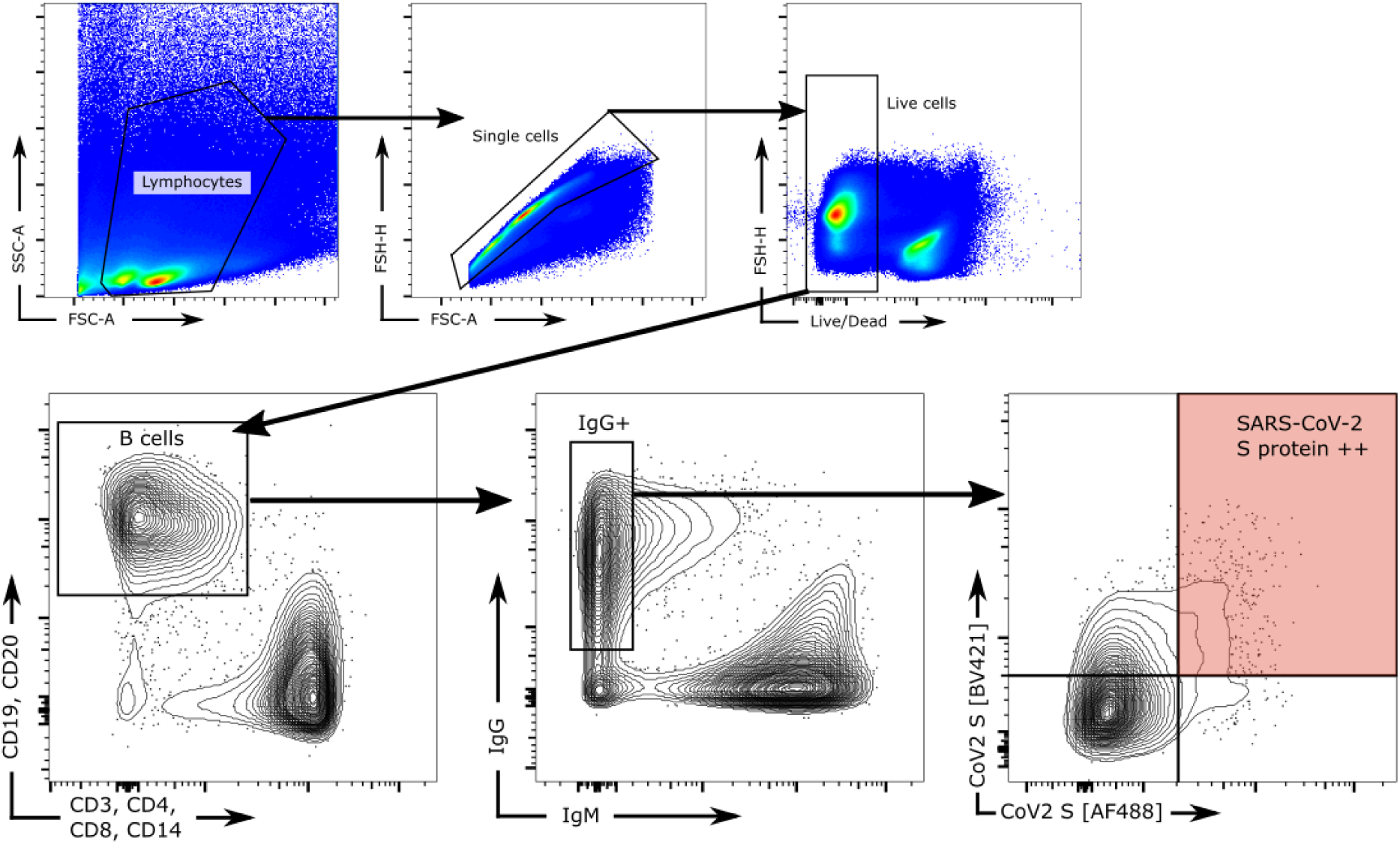
Gating strategy for bulk sorting for 10X Genomics analysis and mAb isolation. Inguinal lymph node samples from all 8 animals were sorted according to this gating strategy. The sorting gate is marked in red. Data were collected during the sort.

**Figure S8.**
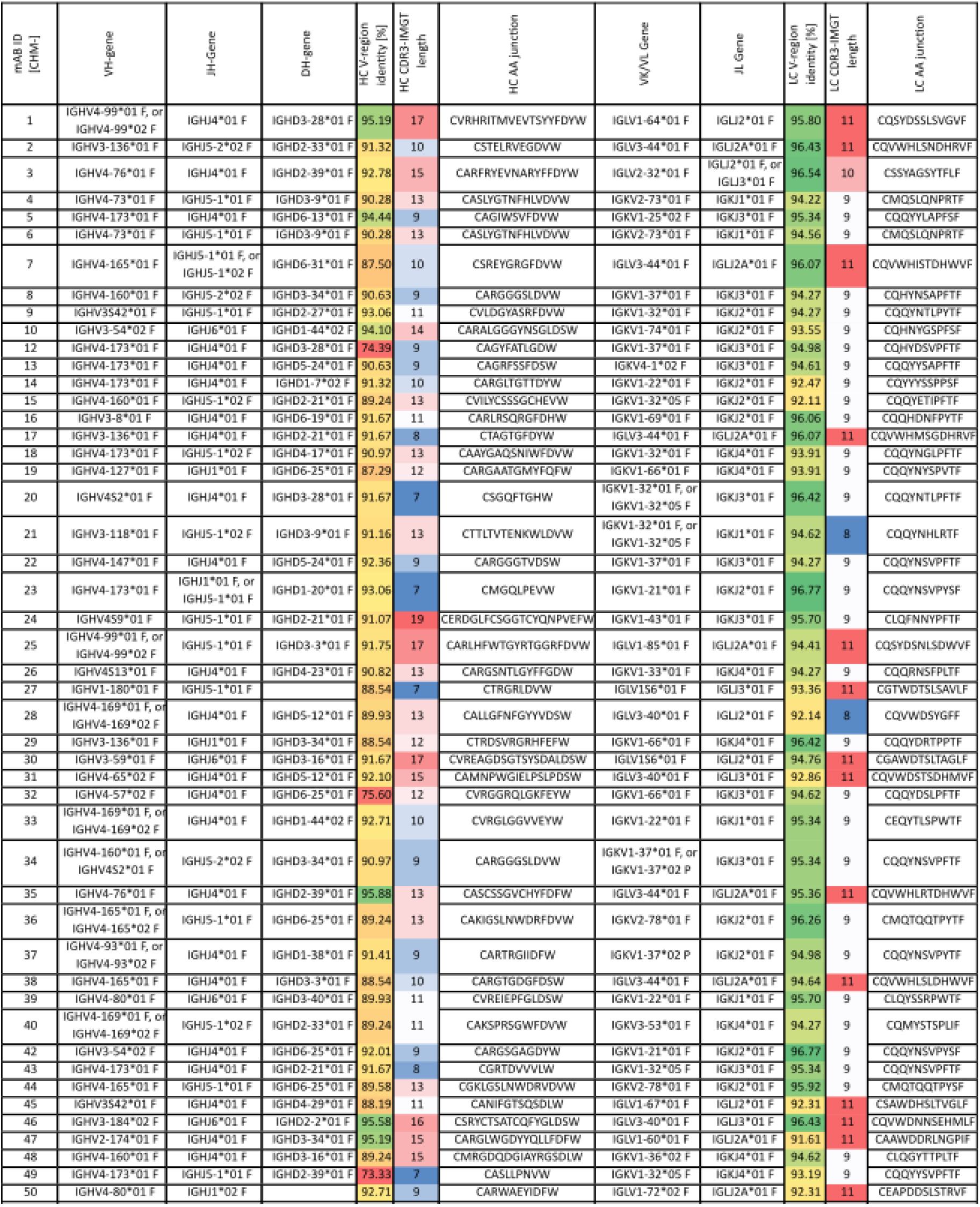
Sequence characteristics of isolated mAbs. The heavy chain (HC) variable region (V_H_) and light chain (LC) variable region (V_K_/V_L_) gene usage, number of V region somatic hypermutations (SHM), complementary determining region3 (CDR3) sequences, and CDR3 lengths, are shown for each mAb. All gene assignment was done using the rhesus macaque (*Macaca mulatta*) germline database from https://www.imgt.org/IMGT_vquest/analysis.

**Figure S9.**
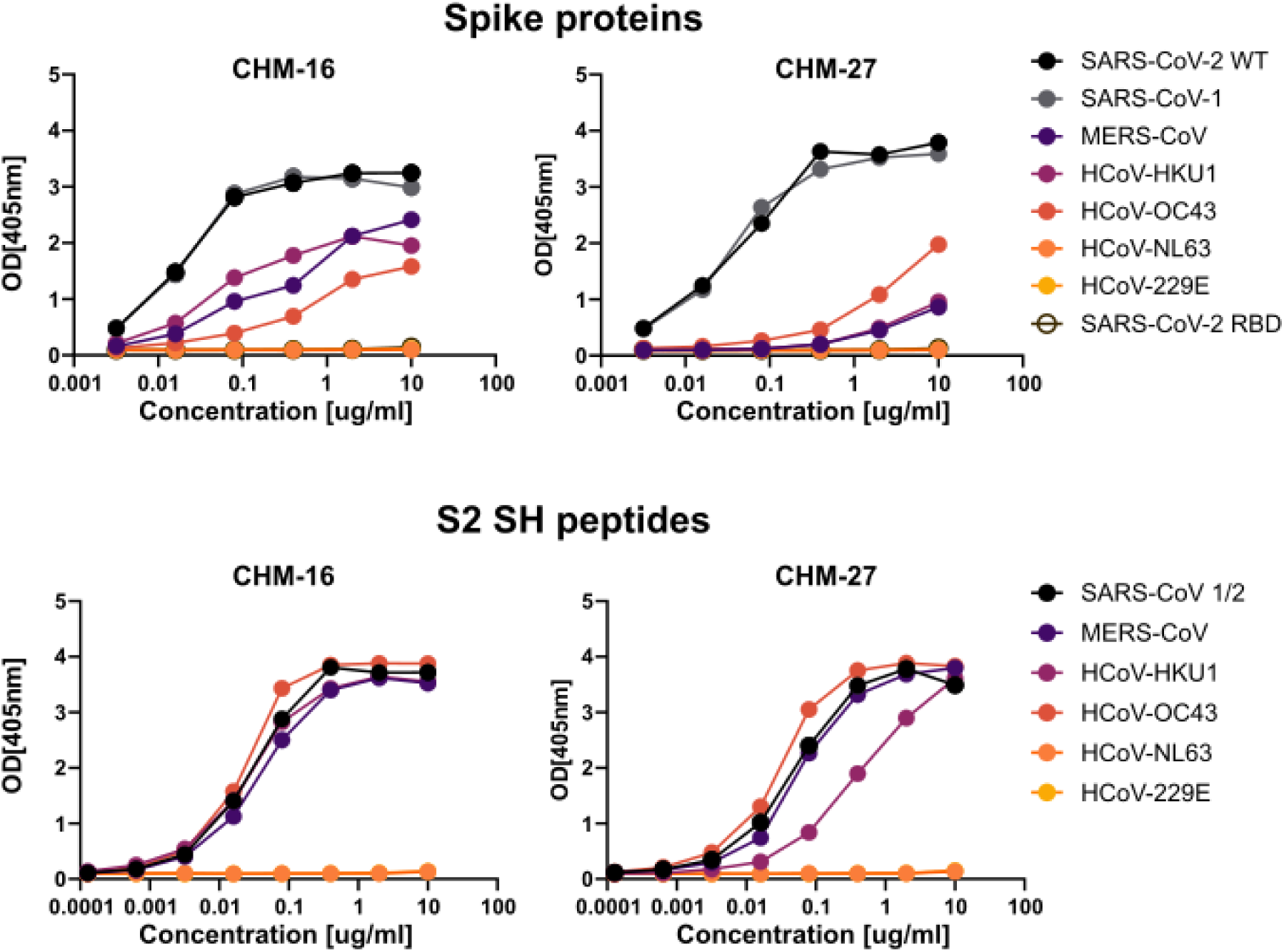
Binding attributes of S2 stem bnAbs CHM-16 and CHM-27. Binding of CHM-16 and CHM-27 to a panel of spike proteins and RBD proteins (top) and S2 stem-helix peptides (bottom) as measured by ELISA.

**Figure S10.**
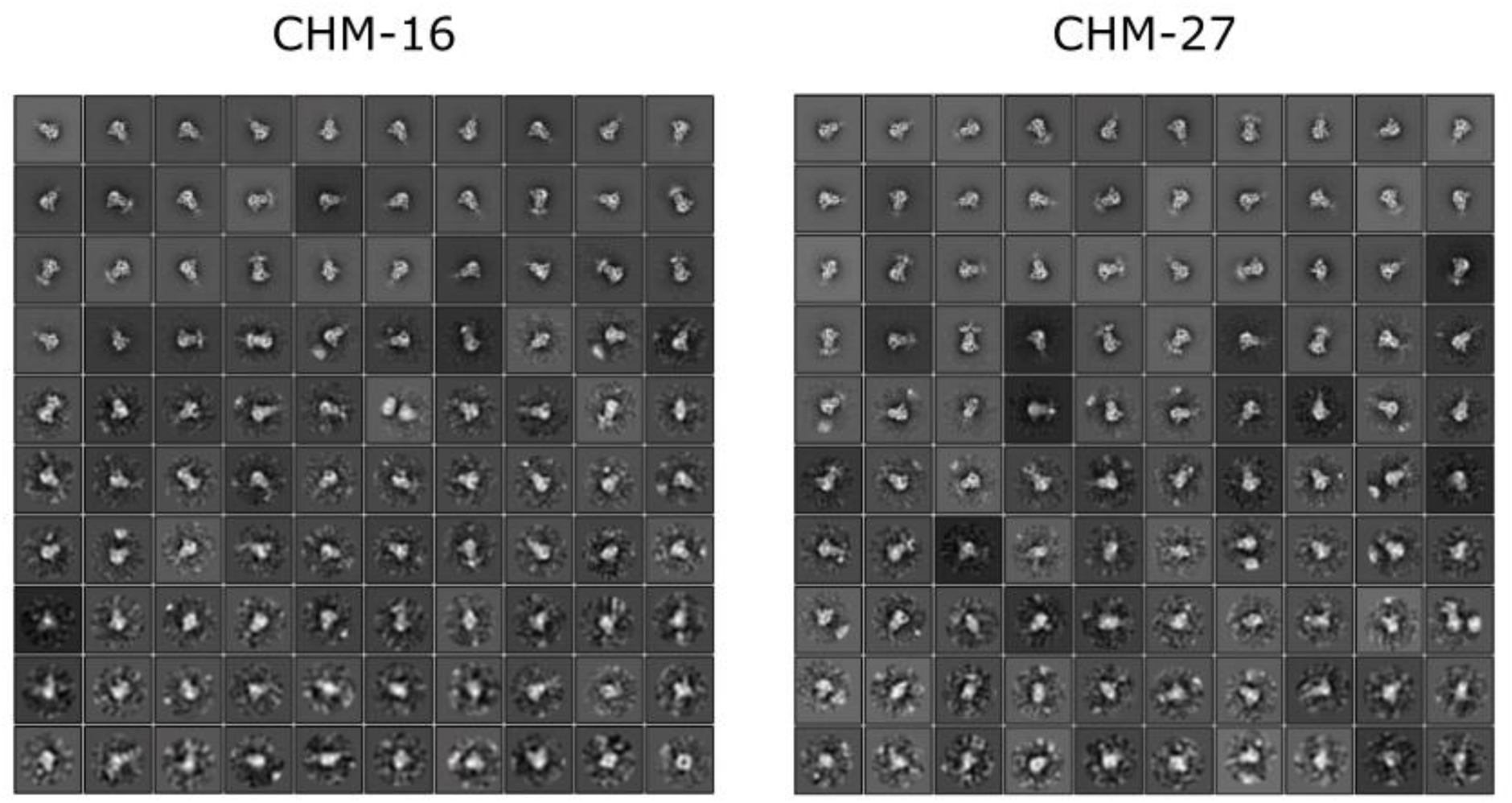
Epitope mapping of S2 stem bnAbs CHM-16 and CHM-27 by negative stain electron microscopy. The 2D class averages of Fabs of CHM-16 (left) and CHM-27 (right) are shown in complex with the SARS-CoV-2 (Wuhan) spike.

**Figure S11.**
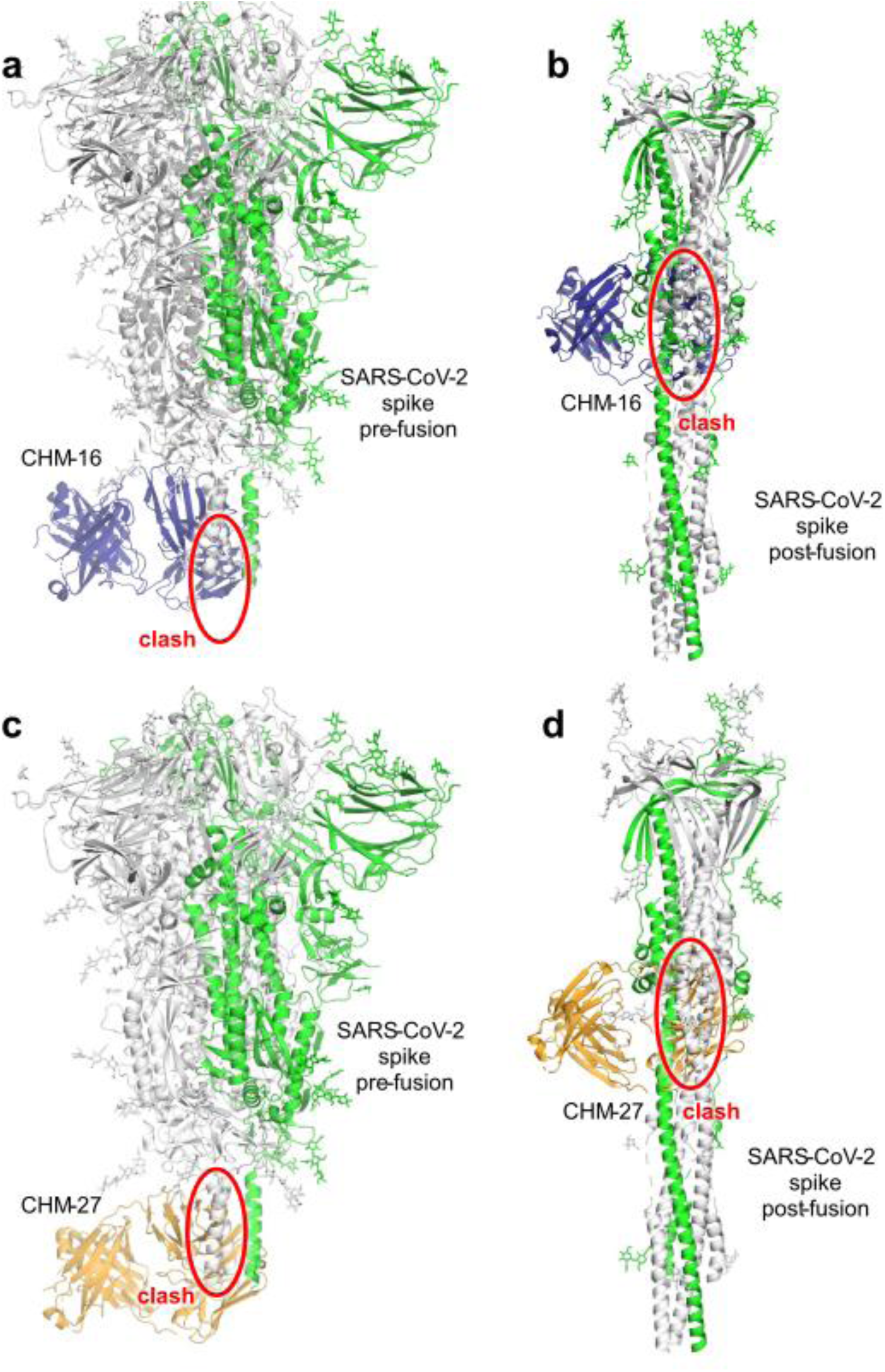
Apparent clashes of mAbs CHM-16 and CHM-27 targeting their S2 stem-helix epitopes on the Spike protein. CHM16 and CHM-27 binding to their S2 stem-helix epitopes would sterically clash with the spike protein in both pre- and post-fusion states and require conformational changes or ‘breathing’ of the epitope region. The crystal structures of CHM-16 in complex with SARS-CoV-2 S2 stem peptide and CHM-27 in complex with MERS-CoV S2 stem peptide were superimposed with SARS-CoV-2 spike structure in its (a,c) pre-fusion state (PDB 6XR8) and (b,d) post-fusion state (6XRA). Superimposed protomers are shown in green while the other two protomers of the spike trimer are in white. Steric clashes are indicated by red circles.

**Figure S12.**
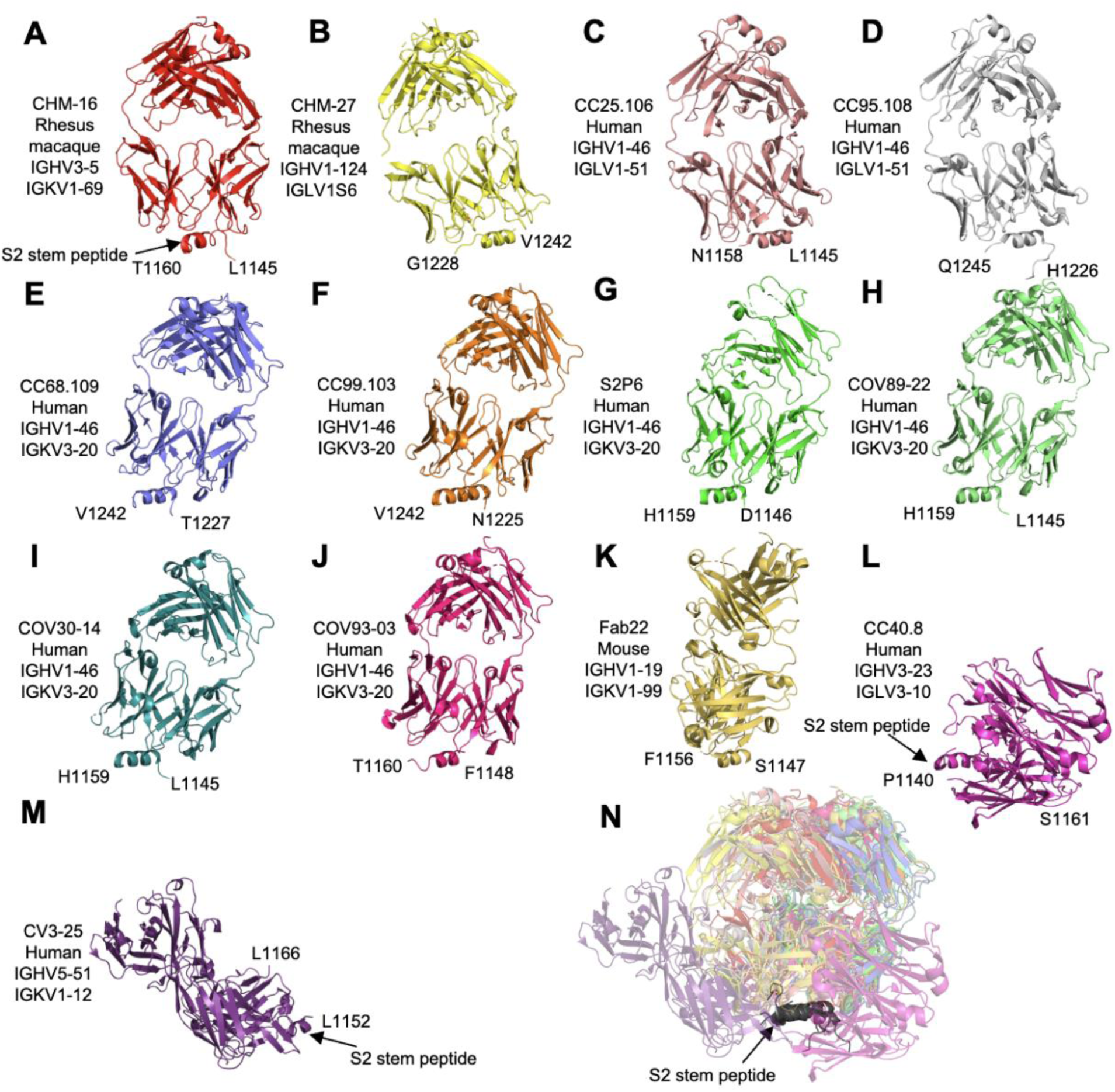
Structural alignment of 13 antibodies targeting CoV S2 stem peptides. All S2 stem helices are shown in the same orientation. Organisms and V_H_/V_L_ germline genes of each antibody are listed. The locations of the S2 stem helices are indicated. (**a**) Structure of CHM-16-peptide complex (this study). (**b-d**) Structures of S2 stem peptides in complex with human CC25.106 (PDB: 8DGU), and CC95.108 (PDB: 8DGW). These antibodies share a nearly identical angle of binding to the stem helix. (**e-j**) Structures of S2 stem peptides in complex with human CC68.109 (PDB: 8DGX), CC99.103 (PDB: 8DGV), S2P6 (PDB: 7RNJ), COV89-22 (PDB: 8DTX), COV30-14 (PDB: 8DTR), and COV93-03 (PDB: 8DTT). These antibodies share a nearly identical angle of binding to the stem helix. (**k**) Structure of mouse Fab22-peptide complex (PDB: 7S3N). (**l**) Structure of human CC40.8-peptide complex (PDB: 7SJS). (**m**) Structure of human CV3-25-peptide complex (PDB: 7RAQ). (**n**) Structural alignment of 13 antibody-peptide complexes superimposed on their S2 stem helices illustrating divergence is the antibody approach to the stem helix. S2 stem helices are shown in gray and antibody colors are the same as panels **a-m**.

**Figure S13.**
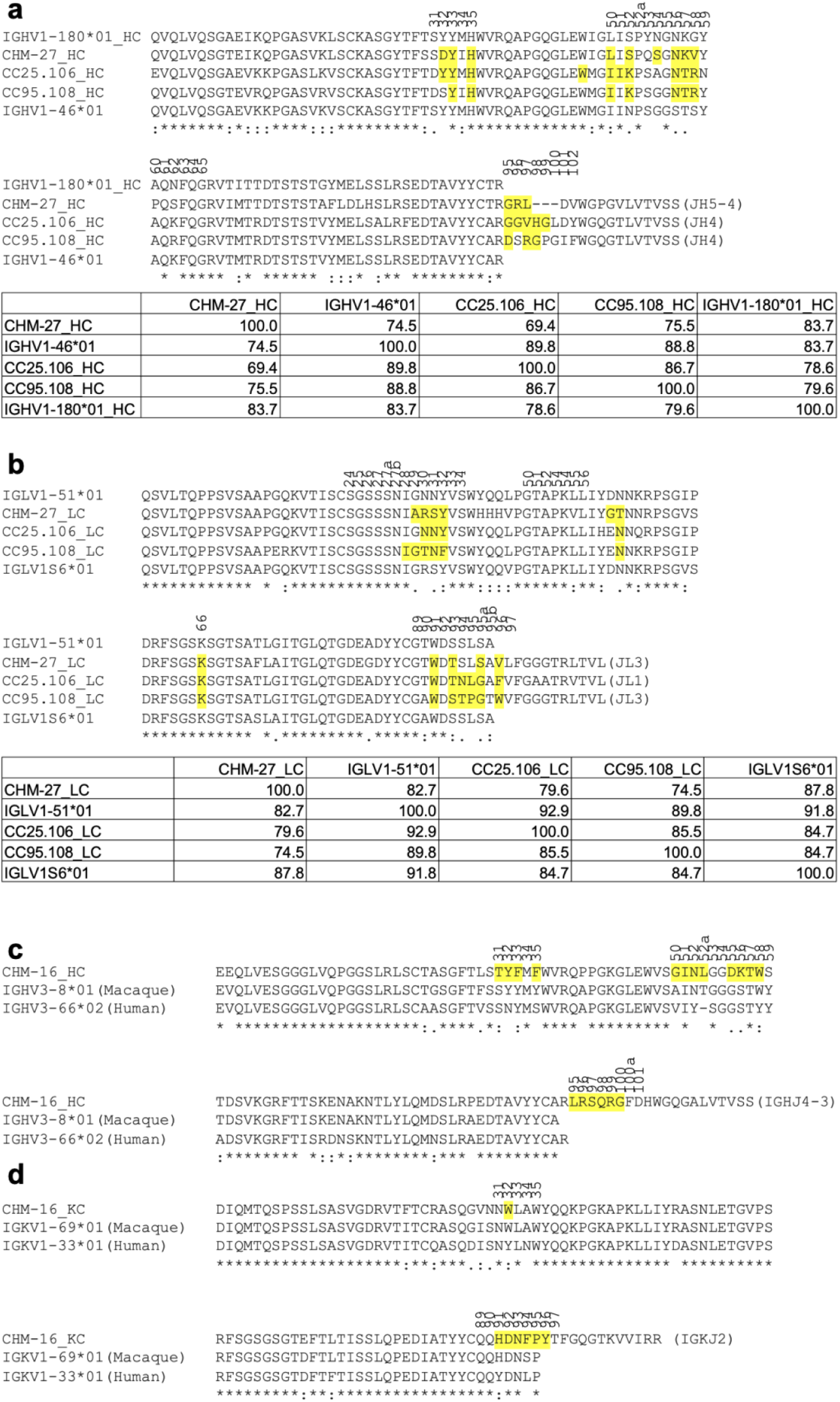
Comparison of rhesus and human S2 stem-helix antibody heavy and light chain sequences. a-b) Sequence alignment of macaque Ab CHM-27 and human Abs CC25.106, CC95.108 with their putative germline genes. Putative germline sequences of (**a**) heavy and (**b**) light chains were conducted by searching the IMGT database(Lefranc, 2003). **c-d)** Sequence alignment of macaque Ab CHM-16 with its putative macaque and closest human germline genes. Putative germline sequences of (**c**) heavy and (**d**) light chains were identified in the IMGT database (Lefranc, 2003). Paratope residues that are involved in interacting with the antigens (BSA > 0 Å^2^ as calculated by PISA)(Krissinel and Henrick, 2007) are highlighted by yellow boxes. The residues in the closest human germline gene (IGHV3-66) that differ from CHM-16 paratope residues are highlighted in red. Residues identical in all aligned sequences are labelled by an asterisk (*), whereas a colon (:) and a period (.) indicate strongly similar and less similar sequences, respectively. The sequence alignment was performed with Clustal Omega (Sievers et al., 2011), where amino acid sequence identity percentages are shown in the tables below the sequences. Residue numbers of CDR loops (Kabat numbering) are shown above the sequences.

**Figure S14.**
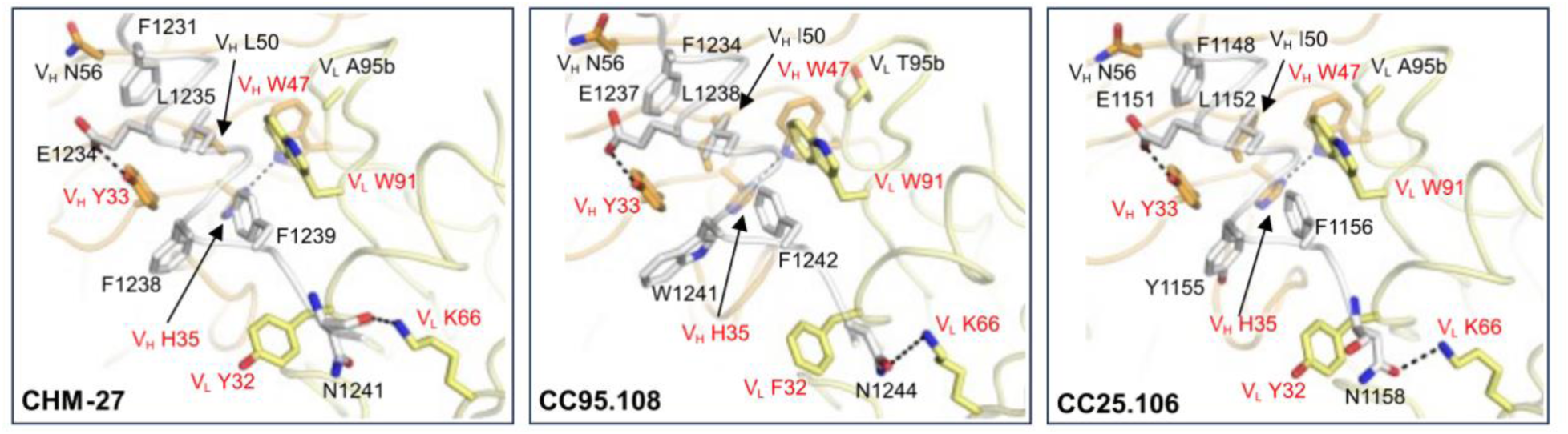
Structural comparison of S2 stem antibodies CHM-27, CC25.106, and CC95.108. Germline-gene encoded residues of CHM-27, CC25.106, and CC95.108 form extensive interactions with the S2 stem helices from MERS-CoV, SARS-CoV-2, and HCoV-HKU1.respectively. Conserved residues that are identical among the three antibodies as well as the macaque germline genes (IGHV1-180 and IGLV1S6), and the human germline genes (IGHV1-46 and IGLV1-51) are highlighted in red. Hydrogen bonds are represented by black dashed lines. As in Figure 6, antibody heavy chain and light chain are shown in orange and yellow, respectively, while the stem helix peptides are depicted in white. Only the variable domains of the Fabs are shown. Kabat numbering is applied to the antibodies. Numbering of the peptides is based on each corresponding virus family.

**Table S1.**
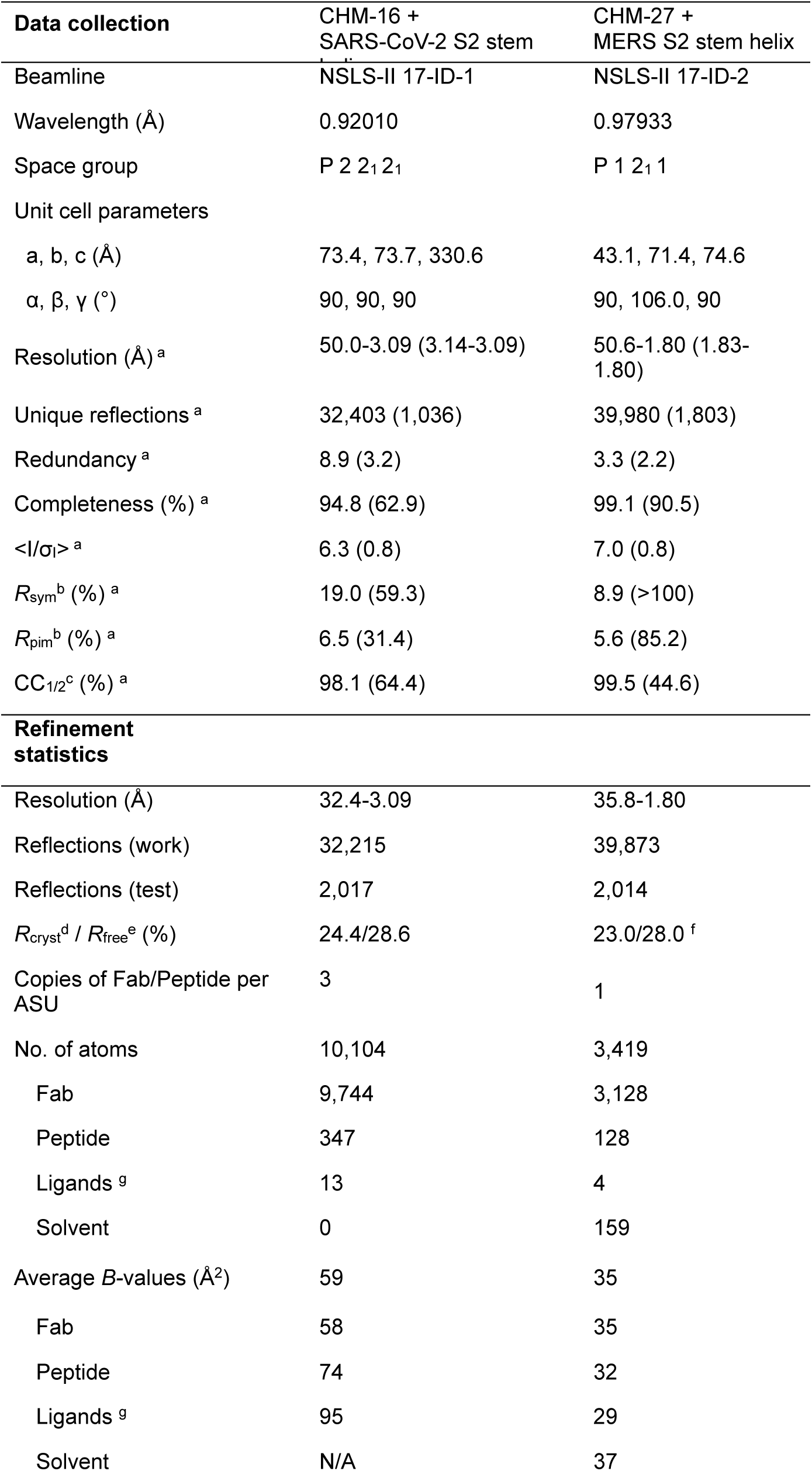
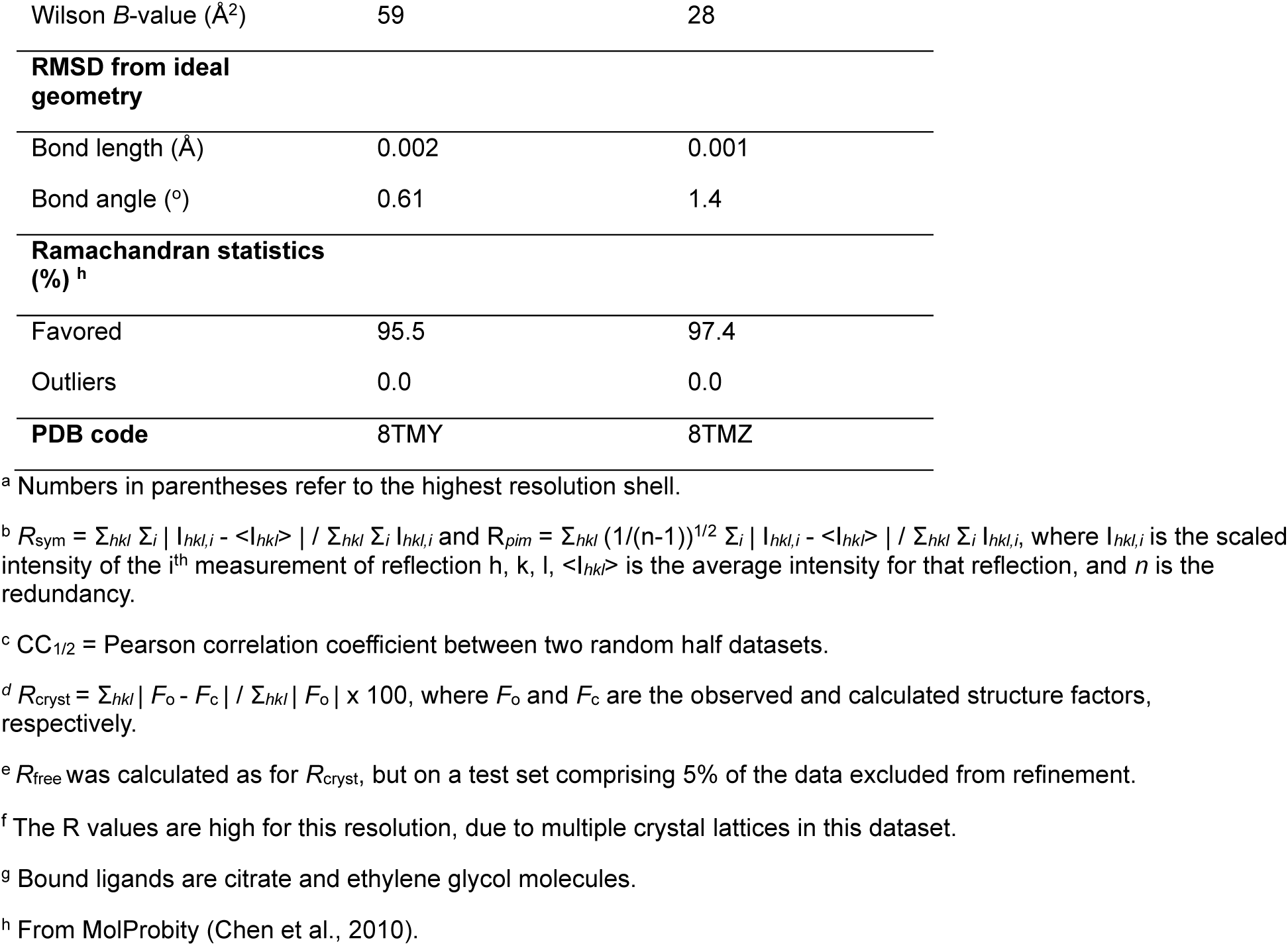
X-ray data collection and refinement statistics.

